# A multiplex, prime editing framework for identifying drug resistance variants at scale

**DOI:** 10.1101/2023.07.27.550902

**Authors:** Florence M. Chardon, Chase C. Suiter, Riza M. Daza, Nahum T. Smith, Phoebe Parrish, Troy McDiarmid, Jean-Benoît Lalanne, Beth Martin, Diego Calderon, Amira Ellison, Alice H. Berger, Jay Shendure, Lea M. Starita

**Affiliations:** Department of Genome Sciences, University of Washington, Seattle, WA, USA; Brotman Baty Institute for Precision Medicine, Seattle, WA, USA; Fred Hutchinson Cancer Research Center, Seattle, WA, USA; Department of Molecular and Human Genetics, Baylor College of Medicine, Houston, TX, USA; Howard Hughes Medical Institute, Seattle, WA, USA; Allen Discovery Center for Cell Lineage Tracing, Seattle, WA, USA

## Abstract

CRISPR-based genome editing has revolutionized functional genomics, enabling screens in which thousands of perturbations of either gene expression or primary genome sequence can be competitively assayed in single experiments. However, for libraries of specific mutations, a challenge of CRISPR-based screening methods such as saturation genome editing is that only one region (*e.g.* one exon) can be studied per experiment. Here we describe prime-SGE (“prime saturation genome editing”), a new framework based on prime editing, in which libraries of specific mutations can be installed into genes throughout the genome and functionally assessed in a single, multiplex experiment. Prime-SGE is based on quantifying the abundance of prime editing guide RNAs (pegRNAs) in the context of a functional selection, rather than quantifying the mutations themselves. We apply prime-SGE to assay thousands of single nucleotide changes in eight oncogenes for their ability to confer drug resistance to three EGFR tyrosine kinase inhibitors. Although currently restricted to positive selection screens by the limited efficiency of prime editing, our strategy opens the door to the possibility of functionally assaying vast numbers of precise mutations at locations throughout the genome.

## Introduction

Resistance to targeted therapies is a major barrier to the successful treatment of many cancers, including non-small cell lung cancer^1–3^. Both *de novo* and acquired resistance to therapeutic small molecules can be caused by mutations in target genes that inhibit drug binding, such as with *EGFR* resistance mutations to covalent EGFR inhibitors. Alternatively, resistance can develop by activating bypass signaling through mutation or amplification of other oncogenes^4^. Pinpointing resistance mutations and developing novel drugs that circumvent them is crucial for continued progress in the treatment of cancer^1^. Mutations associated with resistance can be identified retrospectively in clinical specimens from repeat biopsies or circulating tumor DNA^5–8^. However, obtaining these specimens can be challenging^9^ and causation of resistance must still be verified with functional experiments^10^.

Ideally, resistance mutations would be identified prospectively as part of the clinical development of each drug^1^. Three complementary approaches have been taken to model drug resistance *in vitro*. First, cells can be cultured in the presence of a drug to allow resistant mutations to arise spontaneously, outcompete other cells until they rise to an appreciable frequency, and be identified by whole genome or targeted sequencing^11^. Second, wild type and mutant versions of oncogenes or tumor suppressors can be overexpressed to measure their capacity to provide resistance^12^. Finally, deep mutational scans of open reading frames can identify all potential resistance mutations in candidate genes^10, 13, 14^. However, these approaches are limited in various ways. For example, evolutionary selections are subject to chance, and will likely only identify a subset of potential resistance mutations, while the latter approaches query transgenes rather than mutations in the endogenous genome. An ideal approach would concurrently introduce and characterize vast numbers of potential resistance mutations in genes of interest in their endogenous genomic context. This would allow for upfront identification of resistance mutations to drugs during early stage development, and possible repurposing of agents that can overcome drug resistant variants.

To functionally assess single nucleotide variants or specific small insertions or deletions at scale via CRISPR, a powerful approach is to leverage libraries of homology-directed repair (HDR) templates to install mutations of interest to a short region of interest, *e.g.* an exon. However, the major limitation of this strategy (also known as saturation genome editing (SGE)^15–19)^ is that only one region can be studied per experiment, both because the HDR template library relies on a locus-directing gRNA, and because deciphering which mutation was installed in which cell requires sequencing of the edited locus itself. As such, evaluating candidate resistance mutations in many exons of many genes in response to a panel of compounds via HDR-based saturation genome editing would be highly labor intensive. Saturating scans of multiple exons or genes are possible with base editing, but these lack precision (and range) with respect to which mutations are (or can be) introduced^20, 21^.

Prime editing (PE) is a genome editing method that allows for the precise installation of insertions, deletions, and single base substitutions using a nicking Cas9 fused to a reverse transcriptase (prime editor) and a single prime editing gRNA (pegRNA)^22^. Using PE to assay the effects of single nucleotide variants has two major advantages over other CRISPR-based genome editing methods. First, it allows for multiplex experiments in which any number of mutations in any number of genes can be programmed^23^. This is due to the unique design of the pegRNA, which couples both the target site and programmed edit in a single molecule^22^. Second, this approach takes advantage of the precision of PE, which does not rely on disruptive DNA double strand breaks and exhibits much lower rates of both on-target, intended edits as well as off-target edits, relative to other approaches such as CRISPR/Cas9-mediated HDR or base editing^20, 22^. However, in a recent study deploying PE for multiplex genome editing, the PE outcomes were read out from the edited locus itself^24^. This undercuts the first advantage of PE, as targeting of multiple exons or genes would require each to be serially amplified and sequenced, which increases input requirements and is impractical to scale genome wide.

Here, we present prime-SGE, a scalable, multiplex prime editing framework in which we introduce and assess thousands of precise edits simultaneously in PC-9 lung adenocarcinoma cells. We devised a positive selection strategy to overcome the low editing rate of PE and select for edits that provide resistance to various EGFR inhibitors. In this framework, each cell is engineered to harbor, on average, a single pegRNA encoding a precise edit. This pool of cells is then subjected to tyrosine kinase inhibitor (TKI) drug treatment or a vehicle control. Cells are harvested over a time course of two to three weeks. A key point is that in contrast with other genome editing-based mutational scans^15–20, 23^, prime-SGE achieves readout of the functional selection by quantifying the abundance of prime editing guide RNAs (pegRNAs), rather than the mutated loci. In this proof-of-concept, we found that deep sequencing of the integrated pegRNAs allows for the identification of programmed mutations that give rise to drug resistant cells (**Fig. 1**). Specifically, prime-SGE was able to resolve well-characterized resistance mutations, such as EGFR C797S and KRAS G12C^4^, and also identified potentially novel, previously uncharacterized resistance mutations.

**Figure 1.**
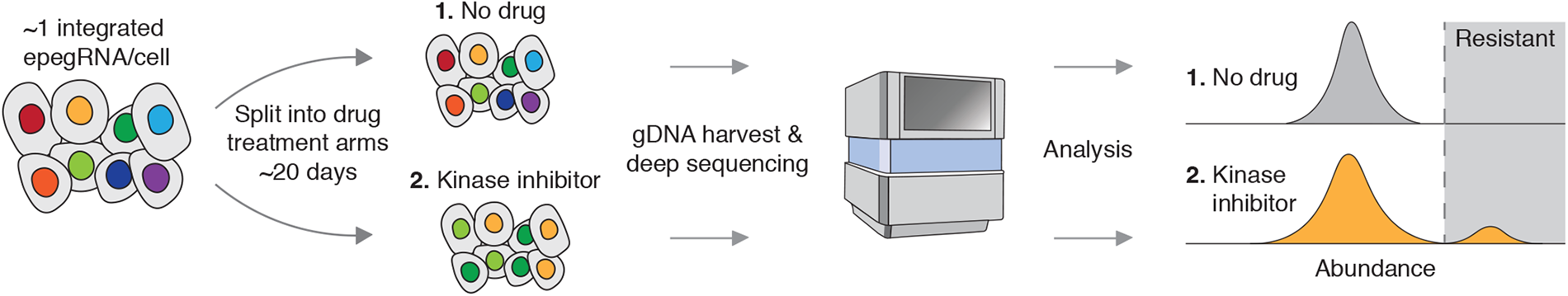
Prime-SGE identifies drug resistance variants at scale. A library of prime editing gRNAs (pegRNAs) are lentivirally transduced into a pool of cells at a low multiplicity of infection (MOI) such that the majority of cells harbor a single pegRNA. This pool of cells is then subjected to treatment with DMSO (no drug) or one of three kinase inhibitors, and cultured for a period of ∼20 days (∼10 cell doublings). Integrated pegRNAs are amplified from genomic DNA and sequenced. Read counts are analyzed to determine the distribution of variant abundances across different treatment conditions.

## Results

### Prime editing of the osimertinib resistance mutation C797S in *EGFR*

As a first experiment, we sought to introduce a single, well-characterized mutation to the *EGFR* gene that confers resistance to the small molecule TKI osimertinib^25^ via prime editing, and then ask whether we could detect its selection during an *in vitro* resistance screen. This mutation, a T to A base change at the first position of amino acid residue 797 of the EGFR open reading frame, changes the wild-type cysteine residue to a mutant serine residue. Osimertinib is a third-generation TKI that targets the ATP binding pocket of EGFR by covalently binding the C797 residue^26^ and is the current standard-of-care therapy for advanced stage EGFR-mutant lung cancer^27^. The cysteine-to-serine change at position 797 blocks the binding of osimertinib and leads to drug resistance and poor survival outcomes^25^. We performed all experiments in PC-9 cells, which are both addicted to EGFR signaling^28^ and sensitive to the TKI osimertinib, providing a model for identification of secondary mutations that confer resistance.

For this initial experiment, we designed three different pegRNAs programming the T to A base change at the first base of residue 797 in *EGFR* (**Table S1**). We performed an arrayed experiment in which we transiently co-transfected plasmids expressing each of these pegRNAs and the PE2 prime editor^22^, into wild type PC-9 cells. Two days later, the cells were treated with osimertinib, and 24 days after drug treatment, cells were harvested and the *EGFR* locus amplified and sequenced (**Fig. 2a**). We observed a 16.8 to 25.3% frequency of T to A edits from reads overlapping the *EGFR* locus across the three pegRNAs tested in contrast with a 0.31% frequency of T to A edits in the wild type, untransfected control, presumably sequencing errors (**Fig. 2b**). As PC-9 cells harbor eight copies of *EGFR,* we estimate that on average, resistant cells had one to two mutated copies of *EGFR* after selection. This experiment confirmed the ability for three different pegRNAs, each encoding the same T to A mutation to 1) successfully program this mutation at a low but appreciable frequency, sufficient for subsequent selection; and 2) confer resistance to osimertinib.

**Figure 2.**
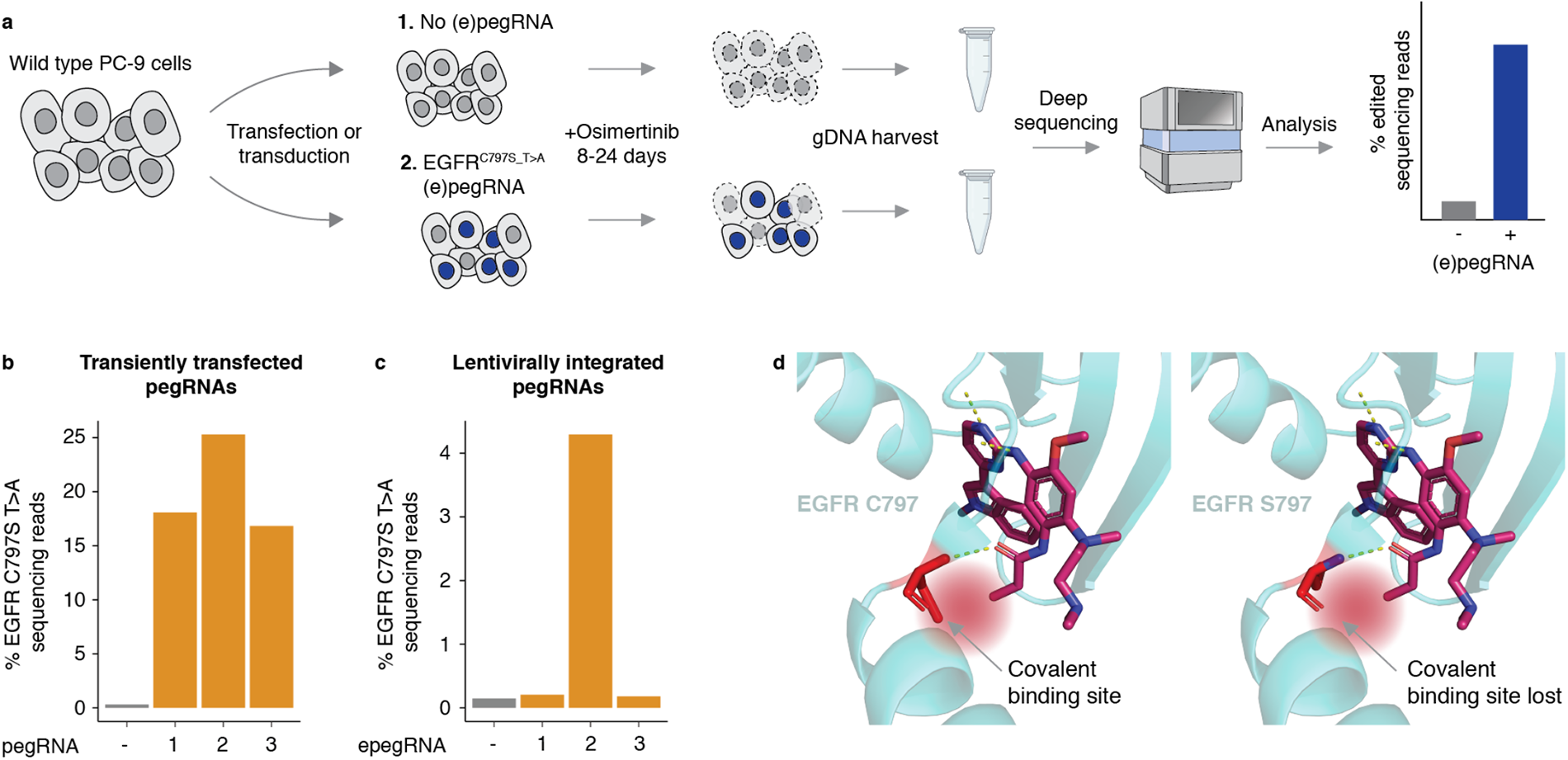
Prime editing of EGFR C797S confers resistance to osimertinib. **a)** Wild type PC-9 cells are transiently transfected or lentivirally transduced with a pegRNA or epegRNA programming the EGFR C797S T>A mutation, along with the prime editor enzyme. Osimertinib is added to the cells for 24 days (transient transfection) or 8 days (lentiviral transduction), genomic DNA is harvested, and the EGFR locus is amplified and sequenced. **b-c)** Percentage of reads with the EGFR C797S T>A mutation after **b)** transient transfection with 3 pegRNAs, or **c)** lentiviral transduction of 3 epegRNAs into PC-9 cells. **d)** Left: Crystal structure of EGFR covalently bound with osimertinib when residue 797 is a cysteine (wild type, Protein Data Bank identifier: 4ZAU)^51^. Right: Crystal structure of EGFR no longer covalently binding osimertinib when residue 797 is mutated to a serine (C797S mutation).

Next, we sought to develop an experimental screening framework in which cells stably express pegRNAs, such that pegRNA identities can be read out by directly sequencing integrated pegRNAs in place of sequencing the edited locus. We modified the LentiGuide-Puro-P2A-EGFP vector^29^ to allow for cloning of pegRNAs (**Fig. S1**). In addition to this lentiviral integration strategy, we also employed the engineered pegRNA (“epegRNA”) construct design that incorporates an RNA stabilizing motif^30^. We also switched from using the PE2 prime editor to using the PEmax prime editor to enable higher rates of editing^31^. We cloned the same three pegRNAs that we used in the transient transfection experiment (**Fig. 2b**) into this modified lentiviral vector, and individually transduced these lentiviral epegRNA constructs into wild-type PC-9 cells. Eight days after osimertinib treatment, cells were harvested and the *EGFR* locus was amplified and sequenced. One of the three virally delivered epegRNAs, which was also the pegRNA that resulted in the highest editing rate in our transient transfection experiment (**Fig. 2b**), successfully edited the EGFR locus and conferred resistance to osimertinib treatment (**Fig. 2c-d**). This experiment confirmed our ability to perform prime editing experiments using integrated epegRNAs, but also highlighted a key challenge, which is that some guides may fail to edit at appreciable frequencies. For all future experiments, we designed up to four epegRNAs per intended mutation.

### Multiplex prime editing resolves well-characterized resistance mutations in *BRAF*, *KRAS*, *EGFR*, *RIT1*, *MET,* and *PIK3CA*

Motivated by our ability to prime edit and select for cells harboring the EGFR C797S T>A mutation, we next performed a pilot screen to assess our ability to install mutations in multiple genes for drug resistance in a single pooled experiment, and to then detect these via guide sequencing (**Fig. 3a, b**). We engineered *MLH1*ko-PEmax-PC-9, a PC-9 cell line with an *MLH1* knockout and integrated PEmax to increase the prime editing rate^31^ (**Fig. S4**). We then designed a library of 121 epegRNAs (**Table S3**, **Table S4**) programming 35 mutations in six oncogenes (*EGFR*, *KRAS*, *PIK3CA*, *RIT1*, *BRAF*, and *MET*, **Fig. 3a**). Nearly all of these mutations have been previously hypothesized to confer resistance to osimertinib, except for the EGFR T790M “gatekeeper” mutation, which was included as a control as it is a known sensitive mutation^26^. We transduced *MLH1*ko-PEmax-PC-9 cells with this epegRNA library. After selecting for cells containing an integrated epegRNA with puromycin, we treated cells with osimertinib. We harvested cells at 21 and 26 days after the initiation of drug treatment. We sequenced both the integrated epegRNA lentiviral construct, as well as the endogenous loci for 31 of 35 programmed edits (**Table S2**), to determine whether the frequency of sequenced epegRNAs matched the frequency of endogenous edits. This dual sequencing approach confirmed that the edits programmed by the various epegRNAs were creating the intended edits in the genome in the untreated (DMSO) arm for 23 of the 31 mutations programmed for which we successfully amplified the endogenous locus, albeit at a low editing rate (between 0.1 and 0.9% per mutation). Further, this approach showed that the epegRNAs that were increasing in frequency in the drug screen were also increasing at the endogenous loci for eight of the 23 successfully introduced edits (**Fig. 3c, d**), suggesting that sequencing the epegRNA can be a proxy for sequencing the edited locus. For the other programmed mutations, we did not observe such consistent increases between the DMSO control vs. osimertinib-treated cells. This is potentially due to the fact that some programmed prime edits were unsuccessful (as evidenced by the lack of detectable editing for 12 of the 35 programmed mutations) and/or because not every mutation included in this library may be a bonafide drug resistance mutation. Of note, for four of the programmed mutations (PIK3CA E453K G>A, MET 1010 splice site variant G>A, MET 1010 splice site variant T>C, and MET 1010 splice site AGGT deletion), we observed increases in epegRNA frequency between the DMSO control and drug conditions, but we did not successfully amplify these targets via PCR so do not have corroborating locus sequencing data.

**Figure 3.**
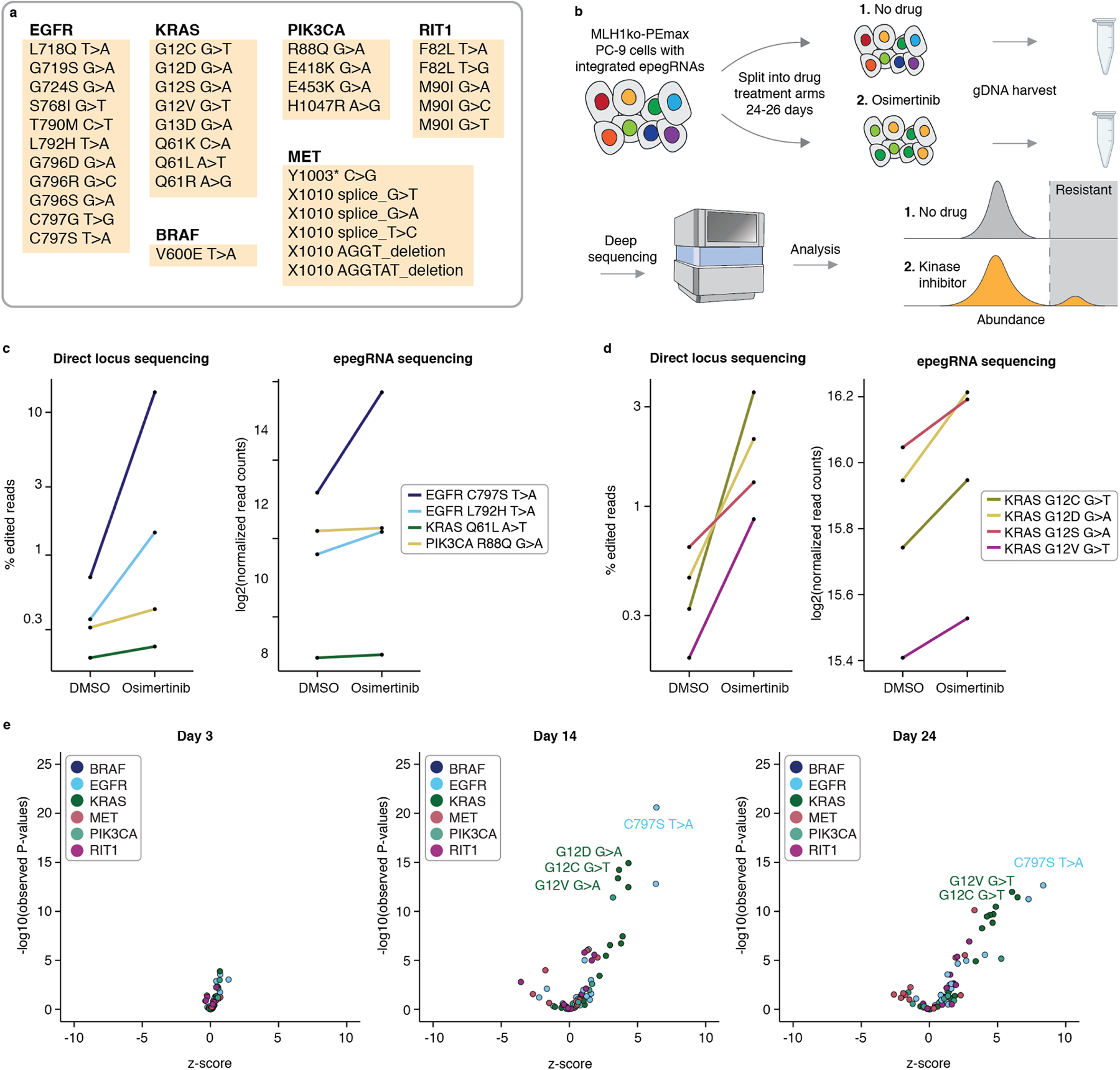
Proof of concept multiplexed prime editing of *EGFR*, *KRAS*, *BRAF*, *PIK3CA*, *MET*, and *RIT1*. **a)** The 35 edits programmed by the 121 epegRNA pooled library. **b)** A pool of 121 epegRNAs was lentivirally integrated into *MLH1*ko-PEmax PC-9 cells and split into a no drug and an osimertinib drug treatment arm for 24 or 26 days. Genomic DNA was harvested and integrated epegRNAs were amplified and sequenced. **c-d)** Left: Percent edited reads by direct locus sequencing results in the no drug (DMSO) and osimertinib drug treatment arms. Right: log2 normalized read counts of the epegRNA that programmed the specified edits. **d)** Same as **c)** but for a different set of targets. **e)** Volcano plots showing the z-scores and p-values (unpaired two-sided t-test) for epegRNAs in the 121 epegRNA pooled library screen at days 3, 14, and 24.

In summary, from this screen, we identified 12 mutations across *EGFR*, *KRAS, PIK3CA* and *MET* that were enriched in the osimertinib-treated cells (log2FC>0 in the drug condition at day 21), including EGFR C797S, EGFR L792H, KRAS G12C, KRAS G12D, KRAS G12S, KRAS G12V, KRAS Q61L, PIK3CA E453K G>A, PIK3CA R88Q G>A, MET 1010 splice site variant G>A, MET 1010 splice site variant T>C, and a MET 1010 splice site AGGT deletion. The EGFR T790M control mutation did not confer resistance to osimertinib, as expected (log2FC=-0.06 in drug condition at day 21).

The results from this screen also exhibited the differential resistance phenotypes of these 12 resistant variants. EGFR C797S is the most well-documented and well-characterized osimertinib resistance mutation, and we observed that it vastly outcompetes all the other identified resistant variants in our screen. From the day 21 to day 26 timepoint, the frequency of the C797S variant rose from 12% to 56% in the epegRNA pool. As such, this differential resistance phenotype indicates that this screening framework can possibly rank variants by their degree of resistance by quantifying the relative fitness of a variant within a pool of cells of many variants, similar to growth-based deep mutational scans^32^ and possibly even reflecting clonal competition that happens during tumor growth.

Using this same library of 121 epegRNAs, we performed a second screen to identify optimal drug concentrations and timepoints to select for edited cells. Cells were treated with three different concentrations of osimertinib (100, 300, and 500 nM), and we harvested cells at 3, 7, 10, 14, 17, 21, and 24 days after drug treatment to profile the rate at which variants were proliferating throughout the timecourse. Replicates were well correlated in this screen across all timepoints (Pearson’s r = 0.73, 0.76, and 0.77 for 100, 300, and 500 nM screens, respectively **Fig. S3b-d**). From this second screen, we identified 18 statistically significant drug resistant variants (log2FC>0, p<0.05, unpaired two-sided t-test between variants and EGFR T790M at days 14-24), including BRAF V600E, EGFR S768I, EGFR G796D, EGFR L792H, EGFR G796R, EGFR C797S, EGFR C797G, KRAS G12C, KRAS G12S, KRAS G12V, KRAS G12D, KRAS Q61L, RIT1 M90I, PIK3CA R88Q, PIK3CA E453K, a MET 1010 splice site G>T variant, a MET 1010 splice site T>C variant, and MET Y1003* early stop codon variant (**Fig. 3e**, **Fig. S3a**, **Table S6)**. As expected, cells with the EGFR T790M mutation did not proliferate in the presence of osimertinib (**Table S6**). We concluded that a 300 nM osimertinib treatment and a timepoint of 10 to 14 days, which corresponds roughly to 7 to 10 doublings (**Fig. S2**), resulted in high signal in this screen (strictly standardized mean difference between EGFR C797S T>A and EGFR T790M = 14.1-14.4). Taken together, the results of these two pilot screens demonstrated that this experimental framework is capable of identifying mutations that confer resistance to TKIs and that sequencing epegRNAs can serve as an effective proxy for sequencing of the edited locus itself (**Fig. 3c-e**).

### Large-scale testing of drug resistance mutations with three inhibitors across seven oncogenes

Osimertinib is the current standard therapy for non-small cell lung cancer patients harboring an EGFR T790M mutation. However, resistance to osimertinib typically develops, on average, within 10 months of treatment due to histological transformation or the acquisition of oncogene amplifications or other resistance mutations^33^. Because of this, newer fourth-generation tyrosine kinase inhibitors have been developed to treat osimertinib-resistant non-small cell lung cancers. Two of these newer inhibitors include sunvozertinib and CH7233163. Sunvozertinib, which is currently in Phase 2 clinical trials and irreversibly and covalently binds EGFR at the C797 residue, is able to treat tumors that have an *EGFR* exon 20 insertion that renders these cells resistant to osimertinib^34^. CH7233163 is a next-generation *EGFR* inhibitor that is a non-covalent, competitive binder of the ATP binding pocket of EGFR^35^ and is currently in preclinical development for treatment of non-small cell lung cancers that are resistant to third-generation inhibitors^36^. Mechanistically, osimertinib and sunvozertinib are similar in that they irreversibly and covalently bind EGFR C797, and CH7233163 differs in that it is a non-covalent, competitive binder of the ATP binding pocket.

Recognizing the unique potential of prime-SGE to scale without the need to sequence each targeted locus, we next asked whether we could apply it to saturate exons and splice site regions of various known oncogenes to screen thousands of single nucleotide variants for resistance to multiple different TKIs in a single experiment. We designed 3,825 epegRNAs programming 1,220 single nucleotide mutations, both missense and synonymous, or deletions in seven different genes (*EGFR*, *KRAS*, *MET*, *RIT1*, *BRAF*, *MEK1*, and *AKT*, **Fig. 4a, b**, **Table S7**, **Table S8**). In designing these epegRNAs, we also included a randomized eight nucleotide barcode directly 3’ of the epegRNA terminator sequence (**Fig. S1e**). The rationale for this barcode was to understand whether resistant cells were clonally derived or whether independent introductions of the same mutation recurrently resulted in resistance.

**Figure 4.**
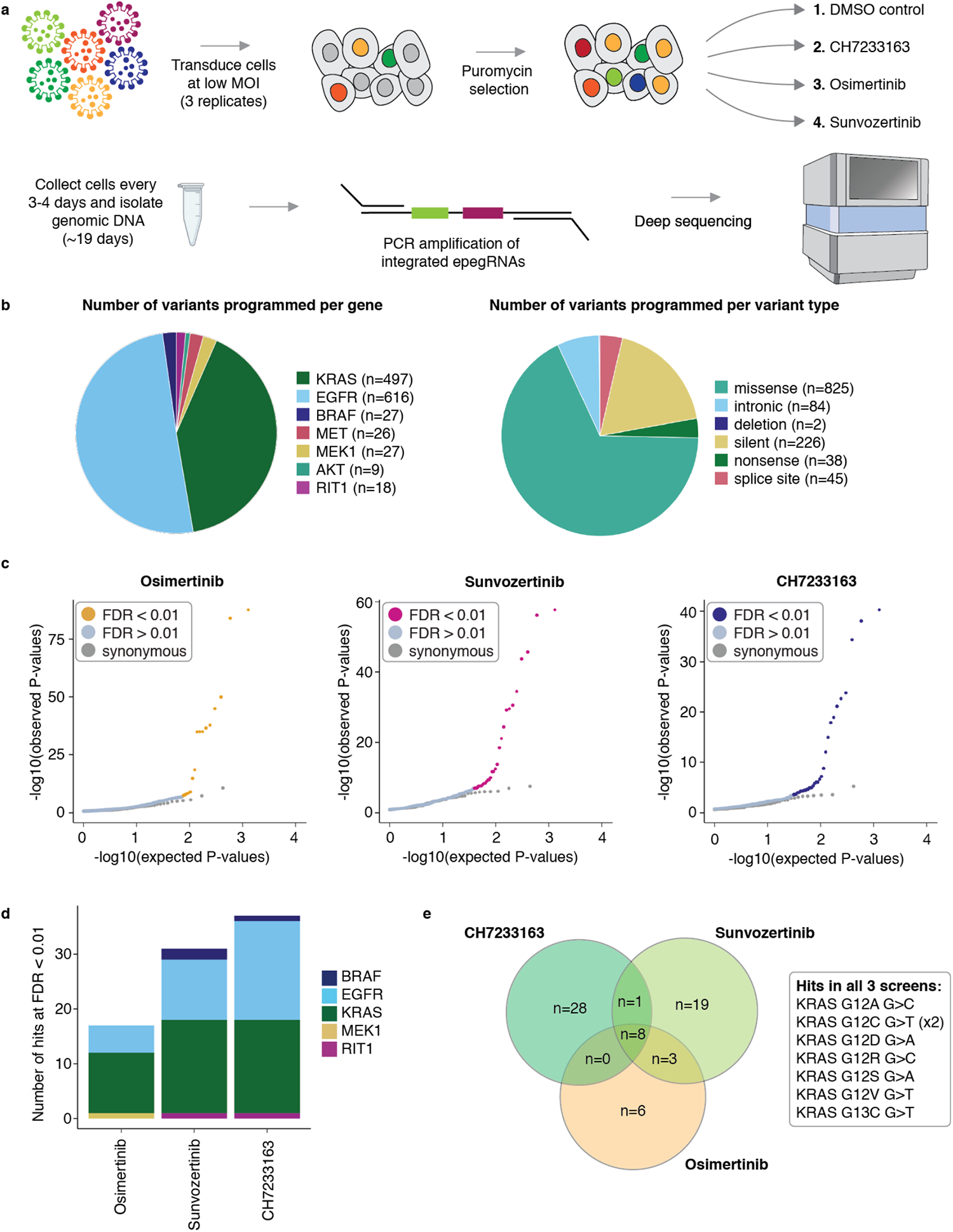
Drug resistance screen of 1,220 mutations against 3 different EGFR inhibitors. **a)** A pool of 3,825 epegRNAs was lentivirally transduced into *MLH1*ko-PEmax PC-9 cells at a low MOI. Cells were selected with puromycin and split into one of four treatment arms (DMSO, CH7233163, osimertinib, and sunvozertinib). Cells were harvested every 3-4 days for over 19 days and integrated epegRNAs were amplified and sequenced from genomic DNA. **b)** Left: Pie chart showing the number of variants programmed for each of seven genes. Right: Pie chart showing the number of variants programmed for each type of mutation. **c)** Quantile-quantile plots showing the distribution of expected versus observed p-values for the three different drug screens. Each point represents a unique genetic variant encoded by 1-4 epegRNAs. **d)** Stacked bar plot showing the number of resistant hits in each screen, colored by gene. Hits are variants that are at an FDR<0.01 and a log2FC>0 by DESeq2 differential epegRNA abundance analysis. **e)** Venn diagram showing the overlap of resistant hits between the three different drug treatment screens.

After generating lentivirus, cells were transduced in triplicate at >2,000X coverage into *MLH1*ko-PEmax-PC-9 cells at an MOI of 0.35 (**Fig. S5**). After puromycin selection, cells were subjected to one of four treatment conditions: DMSO, CH7233163, osimertinib, or sunvozertinib (**Fig. 4a**). Cells were harvested at days 3, 7, 11, 15, and 19, and genomically integrated epegRNAs were amplified and sequenced (**Fig. 4a**). epegRNA read counts were correlated across replicates, timepoints and drug treatments (**Fig. S6**).

We employed DESeq2^37^ to identify hits from this large screen by identifying variants that are differentially abundant between the DMSO control and the three different drug treatments (CH7233163, osimertinib, and sunvozertinib) over the timecourse of 19 days. A likelihood ratio test (LRT) was performed between a reduced model that includes the variable time, and a full model that includes the variables time, drug treatment, and the interaction of drug treatment and time. Synonymous variants were used as controls for false discovery rate (FDR) testing (**Fig. 5b**). From the results of this likelihood ratio test, we identified 17 differentially abundant variants in the osimertinib screen, 31 differentially abundant variants in the sunvozertinib screen, and 37 differentially abundant variants in the CH7233163 screen (FDR<0.01, log2FC>0, **Fig. 4c-e**, **Fig. S7g**).

**Figure 5.**
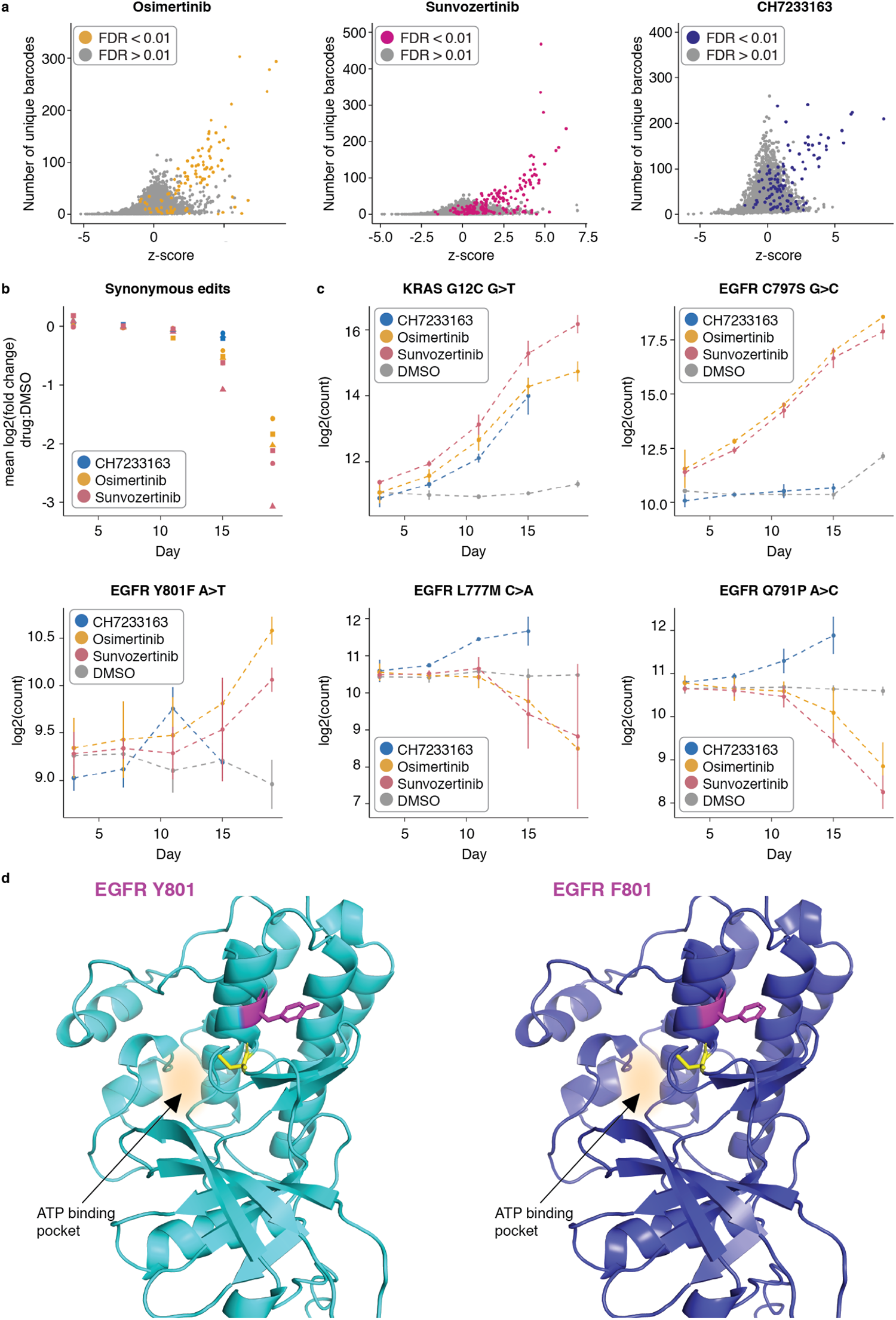
Barcode analysis and individual variant trends across the three scaled drug screens. **a)** Relationship between the z-score and the number of unique barcodes recovered for a given epegRNA. Plots shown include data from two timepoints (day 15 and day 19) and three replicates. Each point represents a unique genetic variant encoded by 1-4 epegRNAs at a single timepoint and a single replicate. Points are colored by FDR (Fig. 4c**). b)** Mean of the log2 fold change of synonymous variants in the three drug screens when compared to the DMSO control. Shapes represent three biological replicates. **c)** Individual variant trends in drug treated cells and DMSO treated cells. Points represent the mean of three replicates, error bars represent one standard deviation from the mean of the three replicates. **d)** Left: AlphaFold predicted structure of EGFR Y801. Right: AlphaFold predicted structure of EGFR F801. Residue 801 is in magenta, C797 residue is in yellow, and the ATP binding pocket is highlighted.

### 3,825 epegRNA screen identifies drug resistance mutations in EGFR and KRAS

There were seven resistant variants that were hits in all three screens, and all seven of these are variants at the 12th and 13th residues of the *KRAS* oncogene (**Fig. 4e**). The *KRAS* gene, which is part of the RAS family of proteins, is the most frequently mutated gene in cancer. Mutations in *KRAS* are a major driver of lung cancers^38^, particularly the *KRAS* G12C mutation. *KRAS*, until the recent approval of sotorasib^39^, was considered to be undruggable due to four decades of failed drug discovery attempts to target this oncogene^38^. Mutations at the 12th and 13th residues in the phosphate-binding loop of *KRAS* are known oncogenic drivers as these mutations directly impair the ability of KRAS to hydrolyze guanosine triphosphate (GTP). This causes the protein to remain in an active GTP-bound state, leading to continued cell growth and the development of cancer. Mutations in *KRAS* do not cause drug resistance by directly inhibiting a drug from binding its target site, but rather by re-activating oncogenic signaling pathways that lead to constitutive signaling and cell growth. All programmed G12 missense mutations were hits in all three screens (p<2.27×10^-7^, LRT), but only a single G13 (G13C) mutation was a hit in all three screens (p<1.58×10^-9^ in all three screens, LRT). KRAS G13D is a known oncogenic mutation^40^, suggesting that the KRAS G13D variant may not have been successfully installed into the genomic DNA via prime editing in this screen.

Another variant which we would expect to give rise to resistant cells in the osimertinib and sunvozertinib screens, but not in the CH7233163 screen, is EGFR C797S, and this is exactly what we observe (**Fig. 5c**). EGFR C797S was a strong hit in the osimertinib and sunvozertinib screens (p=1.14×10^-84^ and p=2.28×10^-55^, respectively; LRT), whereas it was not a hit in the CH7233163 screen (p=0.34, LRT) (**Fig. 5c**). CH7233163 overcomes the EGFR L858R/T790M/C797S triple mutation^35^, and we observe that EGFR C797S was not a hit when cells are treated with CH7233163, suggesting that EGFR C797S does not confer resistance to cells against this inhibitor. CH7233163 differs from both osimertinib and sunvozertinib in that it is not a covalent binder to the EGFR ATP binding site, but rather, is a noncovalent ATP-competitive inhibitor of EGFR. This difference in binding mechanism could explain the differential resistance profiles we observe in cells treated with this inhibitor, such as the lack of EGFR C797S as an identified resistance mutation. Because EGFR C797S is well-edited in this and previous experiments, and because cells in the different drug arms came from the same starting pool of edited cells, it is unlikely to be a false negative in the CH7233163 treatment arm.

In addition to identifying well-characterized resistance mutations, we also identified numerous missense mutations that confer resistance to TKI treatment that are less well-characterized. Examples of such mutations include EGFR Q791P (**Fig. 5c**) and EGFR Q791L, which were both hits in the CH7233163 screen (p=4.35×10^-35^ and p=8.44×10^-13^, respectively). EGFR Q791 missense mutations are not extensively documented as being drivers of TKI resistance, and documented cases of EGFR Q791 in the context of lung cancer, while present, are sparse^41, 42^. Mutations in Q791 are predicted to reduce the binding affinity of EGFR to osimertinib^43^. EGFR Y801 is another example of a residue that when mutated has been identified in single cases of lung cancer^44^, malignant peritoneal mesothelioma^45^, gastric carcinoma^46^ and two cases of squamous cell carcinoma^47, 48^, but it is not known for certain whether a mutation at this residue is a primary driver of resistance to TKIs. EGFR Y801 is a well conserved residue^45^ and lies within the activation loop of EGFR. EGFR Y801F (**Fig. 5c, d**) and Y801N are hits in both the osimertinib and sunvozertinib screens (p<5.96×10^-8^, **Fig. 5c**) but not in the CH7233163 screen. Although our screening framework is not able to conclusively identify non-resistant variants due to the fact that we cannot be certain that prime edits were made, the fact that these EGFR Y801 variants are resistant in two of the three screens is highly suggestive that EGFR Y801 missense variants are successfully being installed by the prime editing machinery, and that they are likely sensitive to CH7233163. Furthermore, the difference in mechanism of EGFR binding between the two covalent inhibitors (osimertinib and sunvozertinib) and the non-covalent inhibitor (CH7233163) plausibly underlies the difference in resistance mutations we observe between these two classes of inhibitors, such as is the case for EGFR C797S. The ability of our screening method to identify rare resistance mutations makes it useful for identifying unknown resistance mutations that have not yet been documented in cancer sequencing databases.

### Barcoded epegRNAs elucidate clonality of resistant cell populations and their growth trajectories

Two features of prime-SGE have the potential to yield unique insights into the emergence and growth behavior of resistant cells. The first of these is the inclusion of a pegRNA-specific barcode, that is directly 3’ of the epegRNA terminator sequence (**Fig. S1e**). The placement of this barcode does not affect epegRNA binding to its target site, as the barcode is not transcribed, but it allows us to amplify and sequence the barcode identity alongside the epegRNA from genomic DNA. The inclusion of this barcode in the epegRNA design allows us to determine whether resistant cells arose from a single versus multiple editing events (and if multiple, exactly how many). To leverage this feature, we first calculated a z-score for each variant to determine its enrichment in the drug treatment conditions vs. control. Next, we examined the relationship between z-scores and the number of underlying barcodes (**Fig. 5a**). This analysis shows that many resistant variants (as identified by DESeq2) are characterized by a high number of unique barcodes (**Fig. 5a**, **Fig. S7a, c, e**). This suggests that, for the most part, resistant cells arose from multiple independent editing events that introduced the underlying resistance mutation. As DESeq2 did not leverage these guide-embedded barcodes, this provides orthogonal support for the classification of many of these hits as resistant. Finally, although this warrants further investigation, this result also suggests that the low efficiency of prime editing is highly target specific – *i.e.* some targets edit well, whereas others are rarely or not at all edited.

A second feature of prime-SGE experimental design is that it allows us to observe the trajectory of resistant cells at high resolution via multiple harvests over the 19 day timecourse (**Fig. 5b, c**). Consistent trajectories over the timecourse adds further confidence to hits – if a variant is increasing in abundance over the timecourse of 19 days in a drug treatment condition as compared to a DMSO control, we can be more certain it is a bonafide resistance mutation. This is useful in a prime editing screen, as the presence of a pegRNA does not guarantee the presence of the programmed mutation. Particularly when identifying potential novel drug resistance mutations, such as EGFR Y801F, observing a consistent increase in the frequency of this variant over the timecourse gives further support to the identification of this missense variant as a statistically significant hit in this screen (**Fig. 5c**).

## Discussion

We describe prime-SGE, a multiplex, prime editing-based screening framework to identify drug resistant variants. A key feature of prime-SGE is that it installs genomic edits via prime editing, which uses a single prime editing gRNA to both encode the target site and the programmed edit. The identity of the pegRNA, and its unique barcode, is read out by sequencing amplified pegRNAs from genomic DNA, which allows for increased scaling of this method to many variants in many exons in many genes in a single screen. Furthermore, individually barcoded epegRNAs enable discrimination and tracking of independently originating editing events, something that is not possible with current saturation genome editing methods. In applying this method to several oncogenes and several tyrosine kinase inhibitors in a lung adenocarcinoma cell line, we were able to resolve well-characterized resistance mutations and oncogenic driver mutations, such as EGFR C797S and KRAS G12 missense variants, as well as less well-characterized resistance mutations, such as EGFR Q791 and Y801 missense variants. We also observed differential resistance phenotypes between covalent and non-covalent binders of EGFR, providing a means to directly compare the resistance behavior of programmed variants across different classes of inhibitors. Looking forward, prime-SGE holds the potential to screen increasing numbers of genetic variants, throughout the genome, for resistance to any number of inhibitors.

Although prime-SGE is able to resolve several drug resistance mutations, a major caveat of this screening framework is that it is unable to conclusively identify all drug-sensitive variants. Despite considerable effort put into designing effective pegRNAs^30, 49, 50^, for any given programmed edit, there remains a high degree of uncertainty that an edit occurred. For example, the KRAS G13 residue is a known oncogenic driver when mutated. However, in our three drug screens, we only identified a single KRAS G13 missense mutation as a hit (KRAS G13C), despite the fact that five other epegRNAs programming G13 missense mutations were present in the library, one of which is a known oncogenic mutation (KRAS G13D)^40^. This suggests that at least the G13D missense variant was not successfully edited into the genomic DNA of cells, rendering our screening framework unable to identify this variant as resistant. This false negative identification of the KRAS G13D mutation surely extends to the other variants we intended to screen, and suggests that the hits we identified are fewer in number than the true number of resistant variants. From screening 1,220 mutations, we identified 17, 31, and 37 drug resistant hits in the osimertinib, sunvozertinib, and CH7233163 screens, respectively. This represents roughly a 2.3% hit rate. It is challenging to determine the false negative rate of this screen, but we hypothesize that we are missing out on the identification of numerous drug resistant variants, as is exemplified by the identification of only one of two expected KRAS G13 missense drug resistant variants.

The prime editing field is advancing at a rapid pace, and we were able to incorporate numerous improvements that were developed over the course of this work. These include knocking out *MLH1* which has been shown to improve efficiency^31^, utilizing PEmax, a further engineered nCas9-RT, and incorporating an improved enhanced pegRNA structural design that includes a 3’ RNA stabilizing motif^30^. With further improvements in editing efficiency, we expect the potential of prime-SGE to grow. With continued improvements, it is also plausible to use this screening framework as a way to identify drug-sensitive variants, to assay drug resistance behavior to combination therapies, to envision prime editing-based screens being used for the large-scale identification of loss-of-function mutations, as has been recently demonstrated^23^, and other functional screening applications. Looking forward, we also envision prime-SGE being applied in increasingly complex cell types and model systems. Particularly if the efficiency issues are addressed, prime-SGE has the potential to greatly accelerate the functional annotation of the extensive list of coding and non-coding variants that underlie risk for both Mendelian and complex human diseases.

## Acknowledgements

We are grateful to members of the Starita, Berger, and Shendure labs for comments, suggestions, and discussions on this work. We are especially grateful for Junhong Choi’s feedback during the initial proof of concept experiments and for discussions with Cole Trapnell regarding how to properly analyze this data. We are particularly grateful to the Shendure lab gene regulation subgroup for technical advice and deep discussions regarding the development of the prime editing screening method. pSpCas9(BB)-2A-Puro (PX459) V2.0 (Addgene #62988) was a kind gift from the F. Zhang lab at the Massachusetts Institute of Technology. LentiGuide Puro-P2A-EGFP (Addgene plasmid # 137729), which was used as the base vector to create the Lenti-epeg-Puro-P2A-EGFP was a kind gift from the F. Wermeling lab at the Karolinska institute. The pCMV-PE2 plasmid (Addgene #132775), pU6-pegRNA-GG-acceptor (Addgene plasmid #132777), and the pCMV-PEmax plasmid (Addgene #174820) plasmids were a kind gift from the D. Liu lab at Harvard University.

## Funding

This work was supported by the NIH NHGRI grant 5RM1HG010461 to L.S and J.S. C.C.S. was supported by the NIGMS (T32GM136534) and NHLBI (F31HL168982). P.P. was supported by a National Science Foundation (NSF) graduate research fellowship. T.A.M was supported by a Banting Postdoctoral Fellowship from the Natural Sciences and Engineering Research Council of Canada (NSERC). J.B.L is a Fellow of the Damon Runyon Cancer Research Foundation (DRG-2435-21). D.C. was supported by the NHGRI (F32HG011817). J.S. is an Investigator of the Howard Hughes Medical Institute. A.H.B. was supported in part by the American Cancer Society Research Scholar Grant RSG-21-090-01 and NIH NCI grant R37CA252050.

## Author contributions

Conceptualization, F.M.C., A.H.B., J.S. and L.M.S.; Investigation, F.M.C.; Data Curation, F.M.C., C.C.S., R.D., N.T.S., and P.P.; Formal Analysis, F.M.C.; Visualization, F.M.C.; Resources, J.S. and L.M.S.; Supervision, A.H.B., J.S., and L.M.S.; Writing – Original Draft, F.M.C.; Writing – Review & Editing, F.M.C., C.C.S., R.D. N.T.S., B.M., P.P., T.A.M., J.B.L., D.C., A.H.B., J.S., and L.M.S.; Funding Acquisition, J.S. and L.M.S.

## Competing interests

J.S. is a scientific advisory board member, consultant and/or co-founder of Cajal Neuroscience, Guardant Health, Maze Therapeutics, Camp4 Therapeutics, Phase Genomics, Adaptive Biotechnologies, Scale Biosciences, Sixth Street Capital, Pacific Biosciences, and Prime Medicine. All other authors declare no competing interests.

## Data Availability

Raw and processed data are publicly available and are accessible via the following website: https://krishna.gs.washington.edu/content/members/multiplex_PE_screening/public/.

## Supplementary Materials

### Supplementary figures

**Figure S1.**
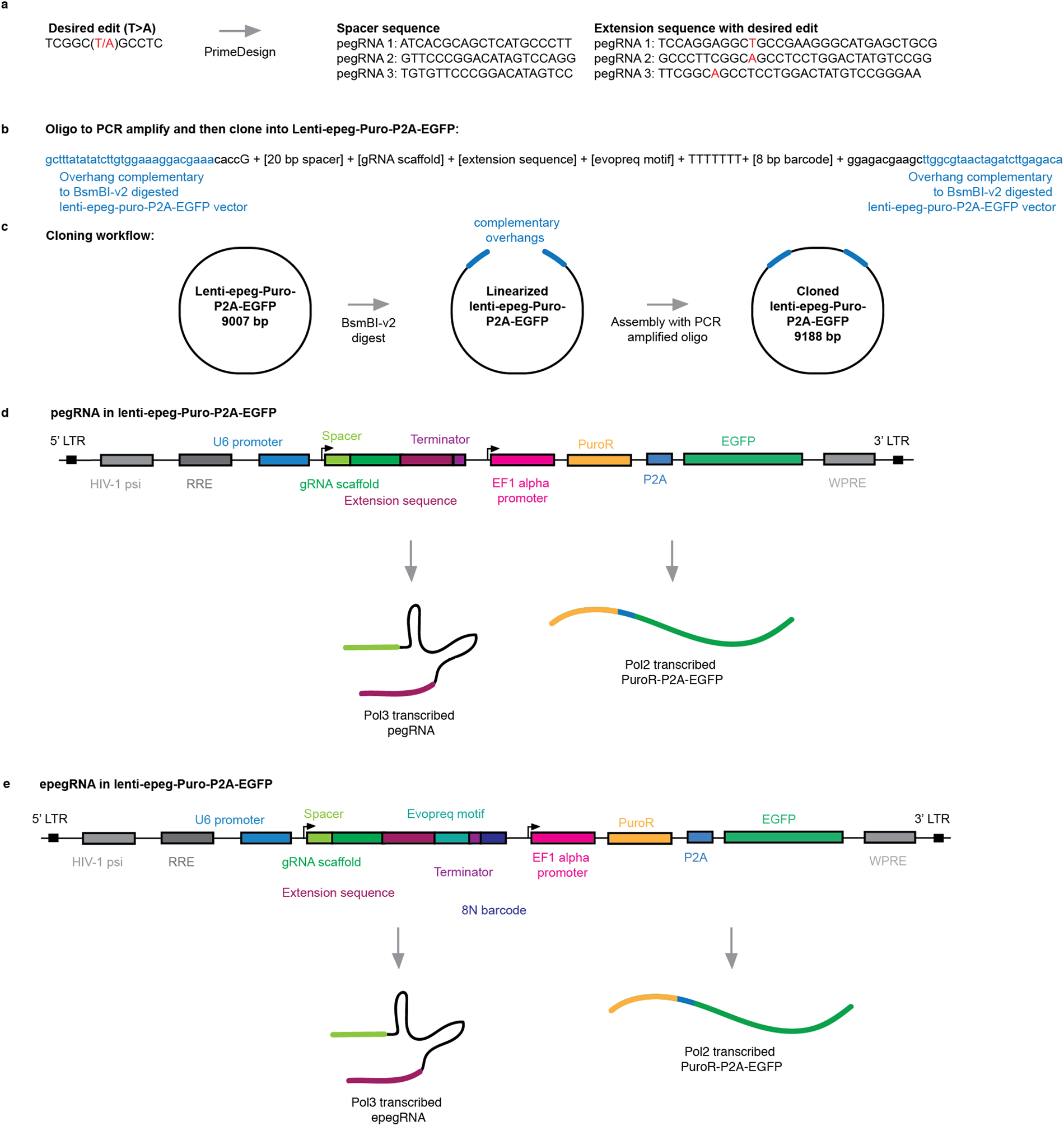
pegRNA design, cloning, and expression vector components. **a)** Schematic of workflow to design pegRNAs with PrimeDesign^49^. Desired edit is in red. **b)** DNA sequence (oligo) to amplify and clone into lenti-epeg-Puro-P2A-EGFP. **c)** Schematic of cloning workflow. Lenti-epeg-Puro-P2A-EGFP is digested with BsmBI-v2, and the PCR amplified oligo from **b)** is assembled with the linearized vector via a Gibson assembly reaction. **d)** Schematic of the lenti-epeg-Puro-P2A-EGFP vector with a pegRNA cloned into it. **e)** Schematic of the lenti-epeg-Puro-P2A-EGFP vector with an epegRNA cloned into it (the epegRNA contains an evopreq RNA stabilizing motif and an 8N barcode sequence 3’ of the terminator sequence).

**Figure S2.**
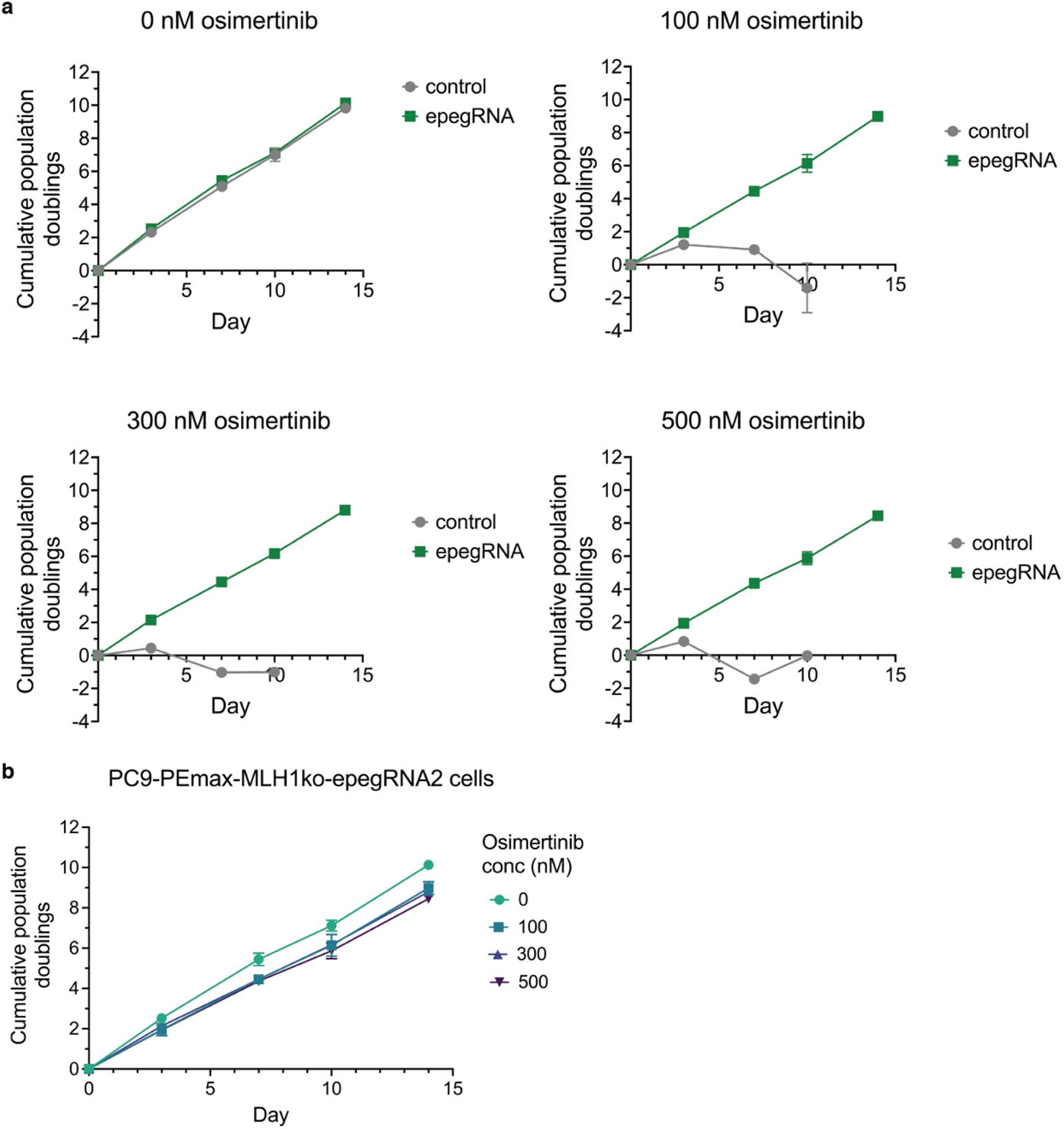
PC-9 cell drug dosing experiments with varying concentrations of osimertinib. **a)** PC-9 cells with no integrated pegRNA (“control”) or a pegRNA programming the EGFR C797S T>A osimertinib resistance mutation (“epegRNA”). Cells were treated with 0, 100, 300, and 500 nM osimertinib and cell population doublings were tracked over a period of 14 days. Data shown are the mean ± standard deviation of two biological replicates. **b)** Same data as in **a)**. Cell population doublings of osimertinib resistant, EGFR C797S T>A harboring cells in 0, 100, 300, and 500 nM osimertinib.

**Figure S3.**
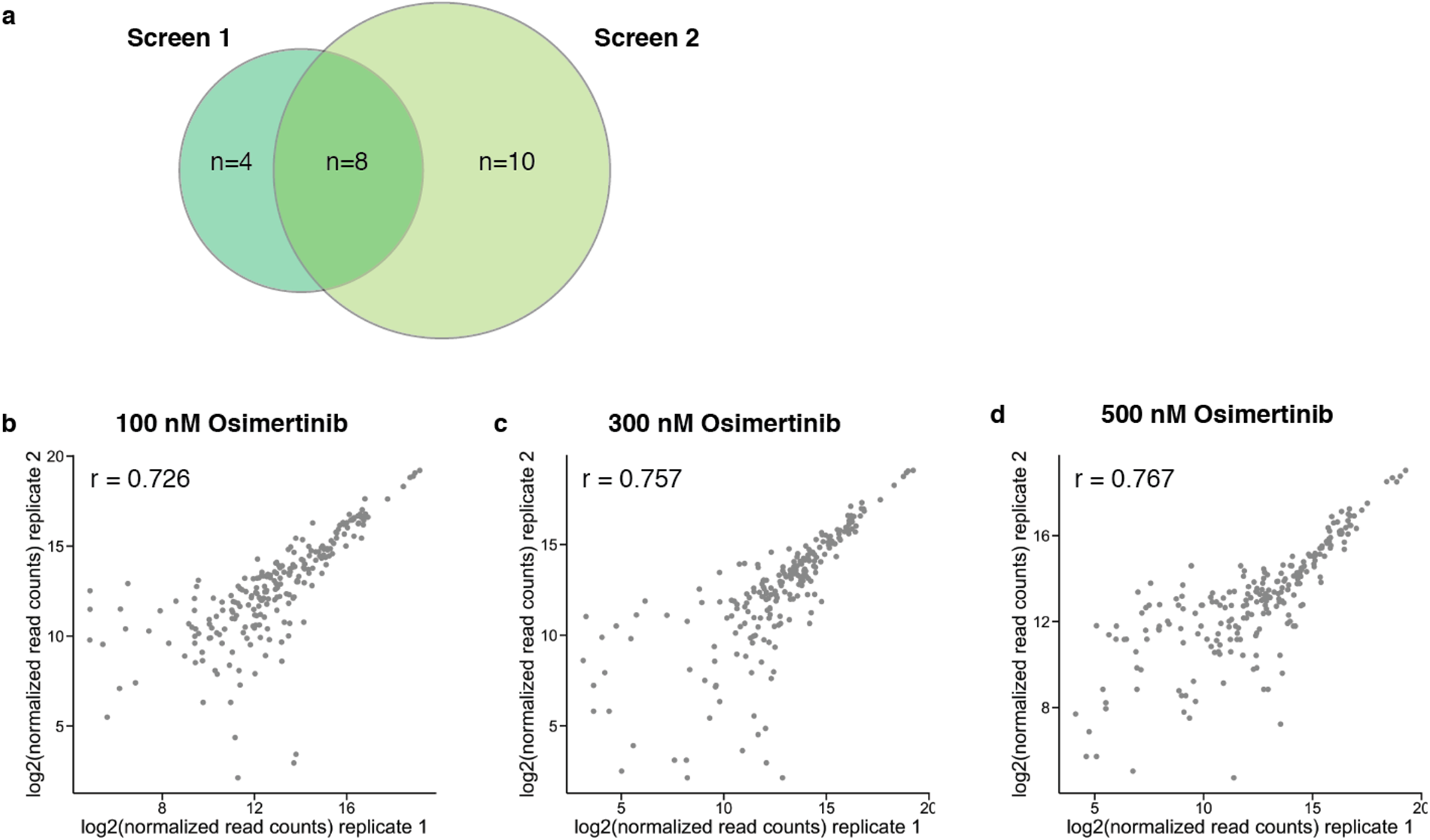
Overlap of resistant variants between 121 epegRNA screens and replicate correlation of second 121 epegRNA screen. **a)** Overlap of hits between the first and second 121 epegRNA screens. **b)** Replicate correlation shown for 121 epegRNA screen at 100 nM osimertinib harvested at seven timepoints (days 3, 7, 10, 14, 17, 21, and 24). Each data point represents a single epegRNA at a single timepoint. **c)** Replicate correlation shown for 121 epegRNA screen at 300 nM osimertinib harvested at seven timepoints (days 3, 7, 10, 14, 17, 21, and 24). Each data point represents a single epegRNA at a single timepoint. **d)** Replicate correlation shown for 121 epegRNA screen at 500 nM osimertinib harvested at seven timepoints (days 3, 7, 10, 14, 17, 21, and 24). Each data point represents a single epegRNA at a single timepoint.

**Figure S4.**
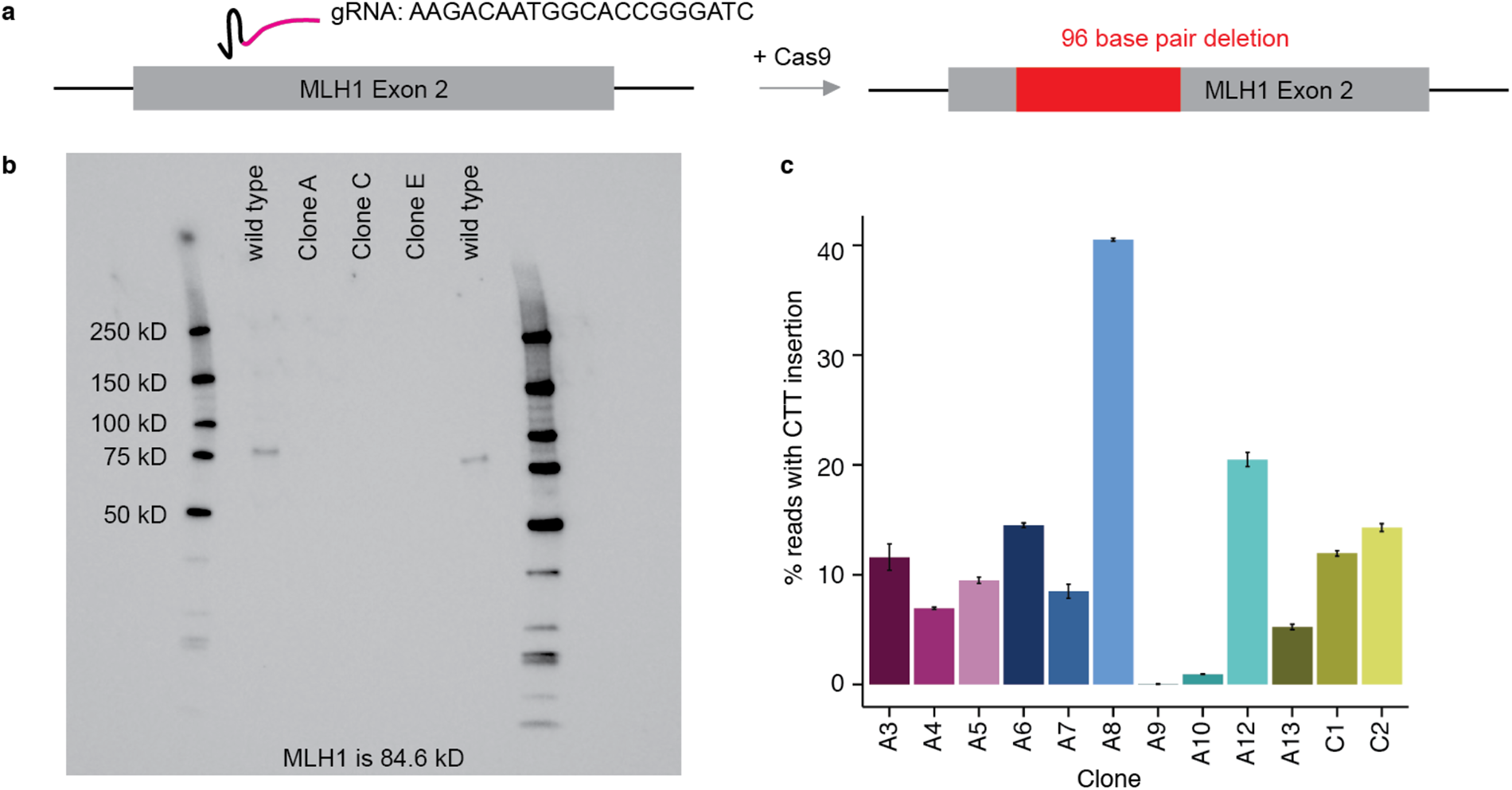
Improvements to prime editing efficiency via an *MLH1* knockout. **a)** A single gRNA (in the pSpCas9(BB)-2A-Puro vector) targeting exon 2 in *MLH1* was transfected into PC-9 cells. This knockout led to a 96 base pair deletion in both copies of *MLH1* in PC-9 cells. **b)** Western blot analysis of three PC-9 knockout clones. All three clones (clones A, C, and E) show complete loss of the MLH1 protein. Wild type cells were run in parallel (lanes 1 and 5) and show presence of the MLH1 protein. **c)** 12 monoclonal *MLH1*ko-PEmax-PC-9 cell lines were tested for insertion efficiency of a trinucleotide CTT insertion at the *HEK3* locus.

**Figure S5.**
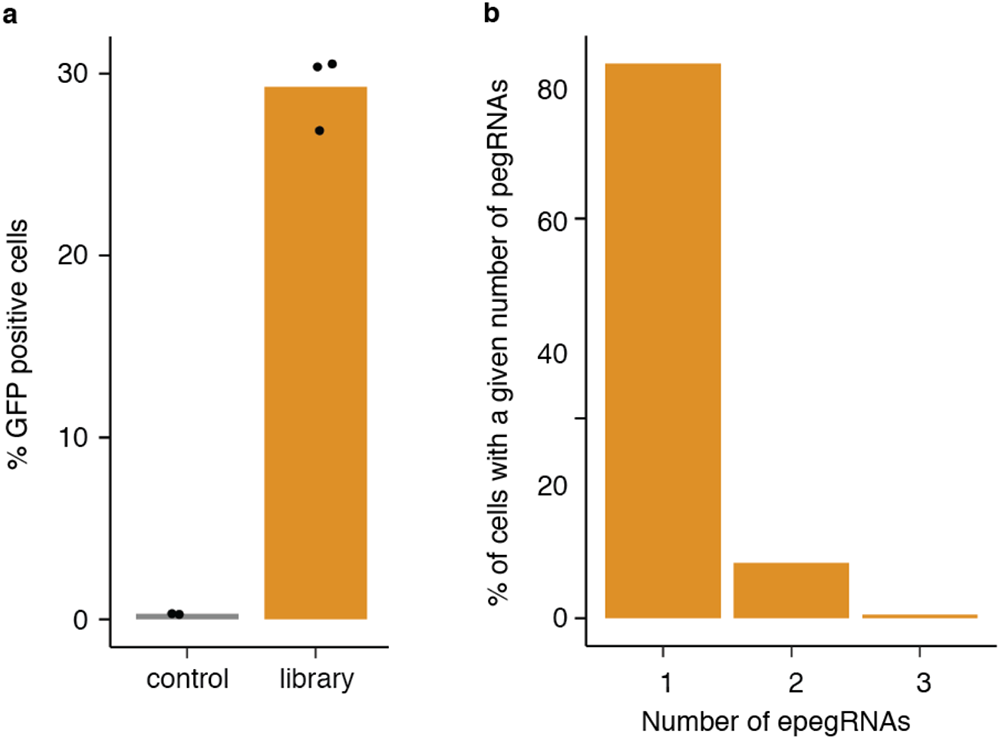
Lentiviral transduction of 3,825 epegRNA library for scaled screen. **a)** *MLH1*ko-PEmax-PC-9 cells were analyzed by fluorescence activated cells sorting (FACS) to analyze the percentage of GFP+ cells following lentiviral transduction of the epegRNA library (GFP is expressed off the lentiviral epegRNA vector). Control cells were not transduced, and the library cells were transduced with the epegRNA library. The three data points represent three independent transduction replicates. **b)** Plot showing the percentage of cells harboring 1, 2, and 3 epegRNAs based off of the achieved MOI (∼0.35) assuming a Poisson distribution of the number of integrations per cell.

**Figure S6.**
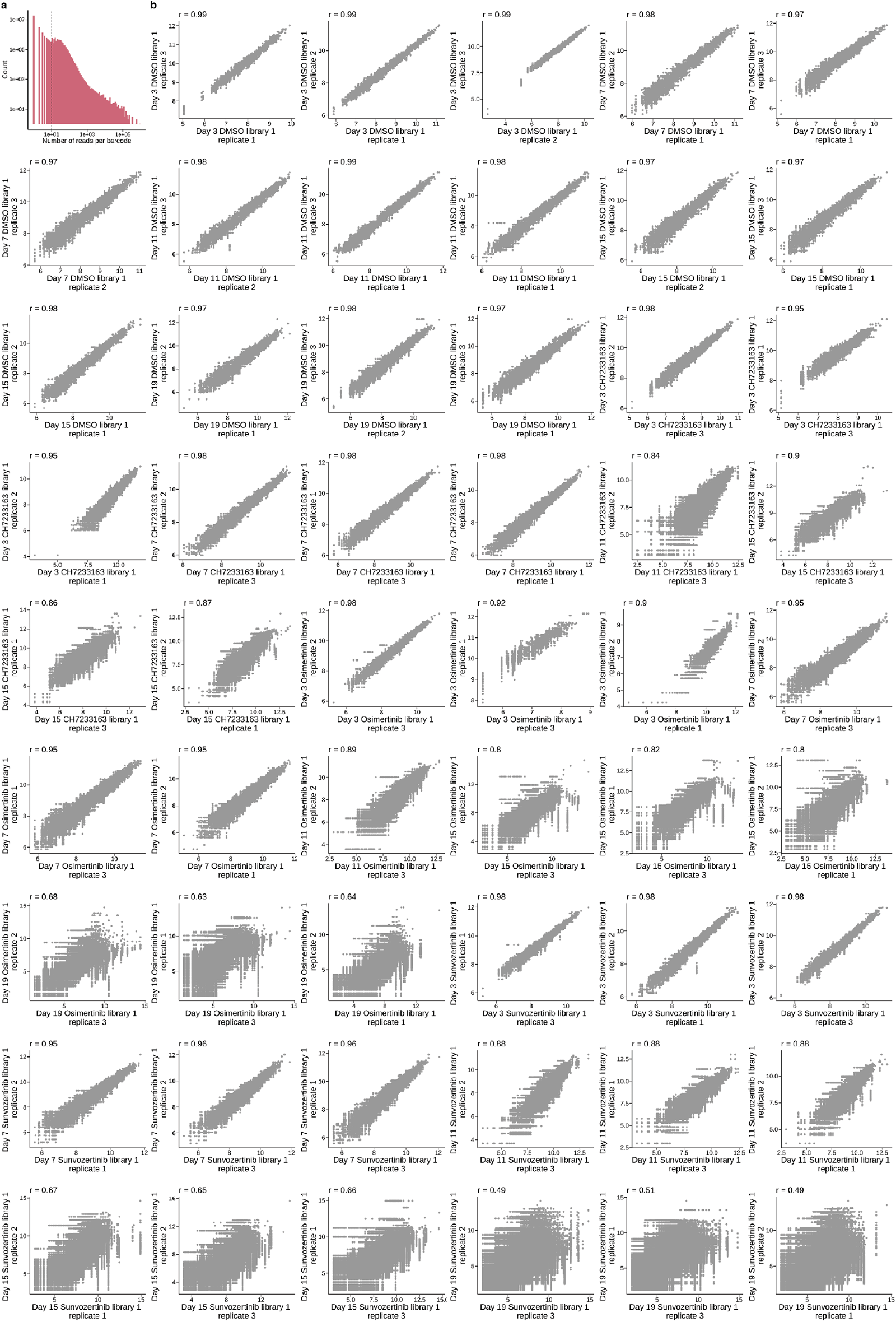
Barcode read count cutoff and replicate correlations in scaled screen. **a)** Histogram showing the number of sequencing reads per barcode. A read cutoff of 10 was used for all analyses. **b)** Replicate correlation plots of log2(normalized read counts) of three independent transduction replicates in the three drug screens.

**Figure S7.**
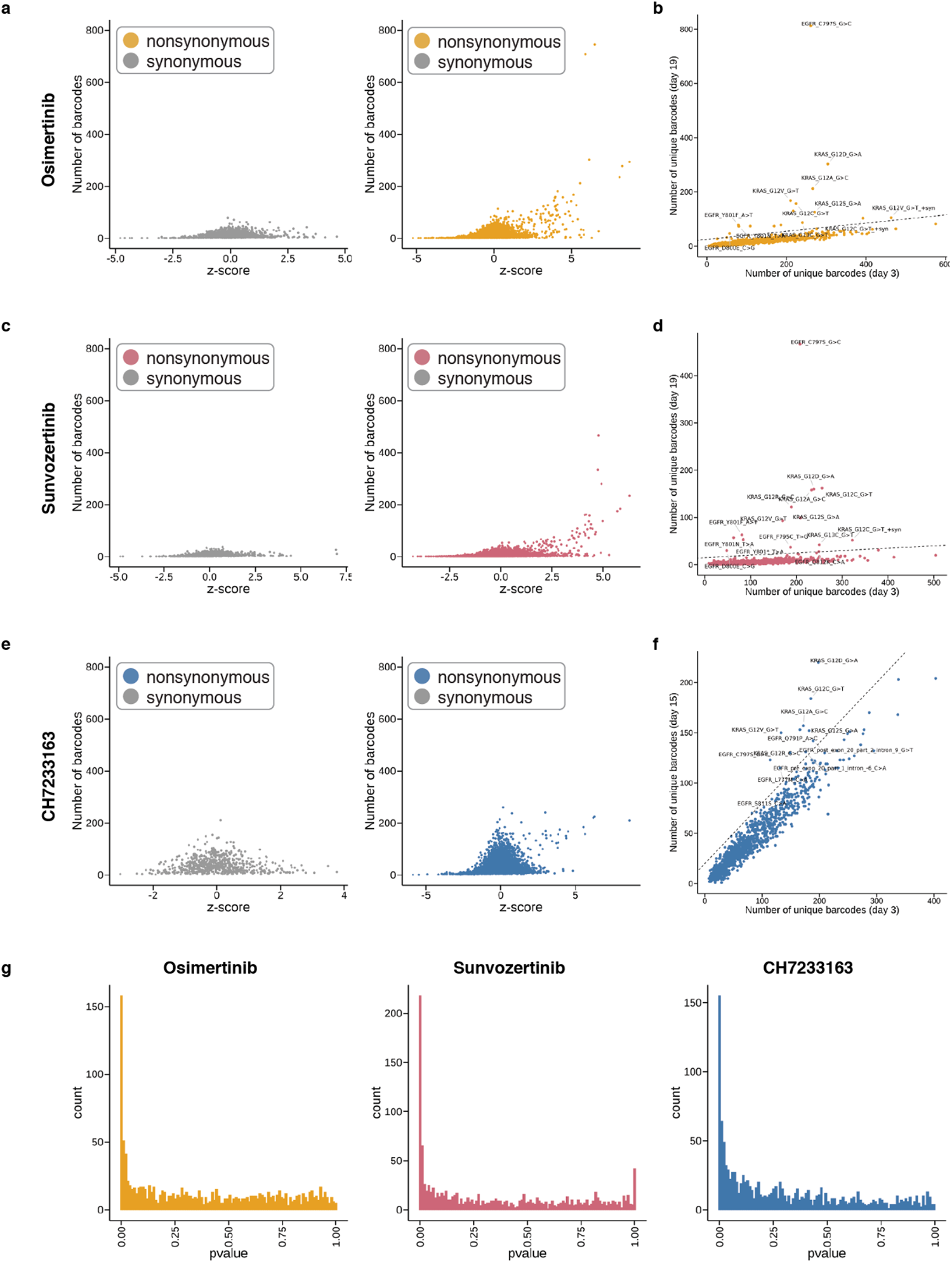
Z-score, barcode count, and p-value statistics from 3,825 epegRNA drug screens. **a)** Z-score and barcode counts plotted for day 15 and day 19 data (combined) for all three replicates for the osimertinib screen. Left: synonymous variants, right: nonsynonymous variants. **b)** Unique barcode count correlation plot between day 3 and day 19 of the osimertinib screen. Variants that fall above the diagonal (y = 0.15x + 25) are labeled. **c)** Z-score and barcode counts plotted for day 15 and day 19 data (combined) for all three replicates for the sunvozertinib screen. Left: synonymous variants, right: nonsynonymous variants. **d)** Unique barcode count correlation plot between day 3 and day 19 of the sunvozertinib screen. Variants that fall above the diagonal (y = 0.05x + 15) are labeled. **e)** Z-score and barcode counts plotted for day 15 and day 19 data (combined) for all three replicates for the CH7233163 screen. Left: synonymous variants, right: nonsynonymous variants. **f)** Unique barcode count correlation plot between day 3 and day 15 of the CH7233163 screen. Variants that fall above the diagonal (y = 0.6x + 20) are labeled. **g)** p-value distributions from DESeq2 differential pegRNA abundance testing for the three drug screens.

### Tables S1-S12

**Table S1** – oligos for EGFR C797S arrayed editing experiments.

**Table S2** – Locus-specific PCR amplification primers.

**Table S3** – Input file used for PrimeDesign pegRNA design tool for 121 epegRNA screen.

**Table S4** – Sequences ordered in Twist oligo pool for small scale screen with 121 epegRNAs.

**Table S5** – epegRNA search sequences for 121 epegRNA pooled screen.

**Table S6** – Statistical testing results from second 121 pegRNA screen harvested at 3, 7, 10, 14, 17, 21, and 24 days.

**Table S7** – Input file used for PrimeDesign pegRNA design tool for 3,825 epegRNA screen.

**Table S8** – Sequences ordered in Twist oligo pool for large scale screens with 3,825 epegRNAs.

**Table S9** – epegRNA library contents and search sequences for 3,825 epegRNA pooled screen.

**Table S10** – DESeq2 likelihood ratio test results for CH7233163 screen.

**Table S11** – DESeq2 likelihood ratio test results for osimertinib screen.

**Table S12** – DESeq2 likelihood ratio test results for sunvozertinib screen.

## Methods

### Cell Lines and Culture

#### PC-9 cell culture

PC-9 cells were originally derived from a metastatic lung adenocarcinoma from a 45 year old male patient. All PC-9 cells were grown at 37°C, and cultured in RPMI 1640 + L-Glutamine (GIBCO, Cat. No. 11-875-093) supplemented with 10% <u>fetal bovine serum</u> (Fisher Scientific, Cat No. SH3039603) and 1% penicillin-streptomycin (Thermo Fisher Scientific, Cat. No. 15070063).

#### Piggybac-PEmax vector cloning

The PB-EFS-PEmax vector was constructed as follows. The PEmax coding sequence from the T7 promoter to downstream of the bGH-PolyA tail was amplified out of the pCMV-PEmax plasmid (Addgene #174820) with the following primers CGCCAGAACACAGGACCGGTTAATACGACTCACTATAGGGAGAG (forward primer) and AGCGATCGCAGATCCTTCGCTAATGTGAGTTAGCTCACTCATT (reverse primer). An inverse PCR amplification was done of the backbone of the PiggyBac-PE2-Blast plasmid (generated by exchanging the puromycin resistance cassette in the Piggybac-PE2-puro plasmid^52^ with a blasticidin resistance cassette) with the following primers: CCCTATAGTGAGTCGTATTAGGTGGCAGCGCTCTAGAACC (forward primer) and GAGTGAGCTAACTCACATTACTTCTGAGGCGGAAAGAACC (reverse primer). These two amplification products were then assembled into a single vector (PB-EFS-PEmax) using NEBuilder^®^ HiFi DNA Assembly Master Mix (New England Biolabs, Cat. No. E2621S) using the standard protocol for a 2-3 fragment assembly. 1 uL of the 20 uL assembly reaction was transformed into 50 uL of stable competent E. coli cells (New England Biolabs, Cat. No. C3040H) using the NEB 5 minute transformation protocol. 100 uL of transformed E. coli cells was plated on an LB agar plate containing ampicillin, and single colonies were picked 1 day later to grow up and extract plasmid DNA using a Monarch Plasmid Miniprep Kit (New England Biolabs, Cat. No. T1010L). Extracted plasmid DNA was sequence confirmed via long-read Nanopore sequencing (Primordium Labs) and DNA from a single clone harboring the correct assembled sequence was used for all experiments.

#### *MLH1* knockout-PEmax cell line generation and validation

##### *MLH1* knockout, selection and single-cell sorting

*MLH1* was knocked out of a population of PC-9 cells using a single gRNA targeting exon 2 of *MLH1*. The sequence of this gRNA is AAGACAATGGCACCGGGATC. This gRNA was cloned into pSpCas9(BB)-2A-Puro (PX459) V2.0 (Addgene #62988) via the Zhang lab protocol (https://media.addgene.org/data/plasmids/62/62988/62988-attachment_KsK1asO9w4owD8K6wp8.pdf). 2.5 ug of assembled vector was transiently transfected into 250,000 wild type PC-9 cells using the transIT-LT1 transfection reagent (Mirius Bio, Cat. No. MIR 2300). 2 days after transfection, 1 ug/mL concentration of puromycin (GIBCO/Thermo Fisher Scientific, Cat. No.<u>A11138</u>03) was added to cells to select for successfully transfected cells over a period of 4 days.

After puromycin selection, this population of cells was single-cell sorted into 96-well plates to grow up clonal cell lines. 12 clonal lines were expanded, split into two sets of parallel cultures (i.e. 24 wells with 2 wells per clonal line), and one set was treated with 1.5 uM 6-TG (Sigma Aldrich, Cat. No. A4882) for 4 days to screen for cells with *MLH1* successfully knocked out. 5 of 12 treated wells survived 6-TG treatment (denoted clones A, B, C, D, and E). PCR primers targeting *MLH1* (forward primer: TGTATGAGCCTGTAAGACAAAGGAA, reverse primer: CATCCATATTGAAGCCTTCCTGAAC were used on extracted gDNA from these 5 clonal lines to amplify the *MLH1* locus and confirm knockout via sanger sequencing. A western blot was performed on 3 of the monoclonal lines to confirm knockout (primary antibody used: MLH1 Monoclonal Antibody; Invitrogen, Cat No. MA5-15431, secondary antibody used: Goat anti-Mouse IgG (H+L) Secondary Antibody, HRP; Invitrogen, Cat. No. 31430), and complete loss of the MLH1 protein was confirmed for two clones (A and E) (**Fig. S4**). Clones A and E were chosen for further cell line engineering. Clone E, was ultimately used for all experiments.

##### Prime Editor-max (PEmax) transfection, selection and single-cell sorting

Clones A and E were transfected with a Piggybac-PEmax plasmid with the Piggybac transposase (System Biosciences, Cat. No. PB210PA-1) at a 10:1 molar ratio using the transIT-LT1 transfection reagent (Mirius Bio, Cat. No. MIR 2300). 2.5 ug of total DNA (Piggybac-PEmax plasmid, and the transposase plasmid), along with 5 uL of trans-IT reagent was reverse-transfected into 100,000 cells for each well of a 12-well plate. Cells were selected with 10 ug/mL of blasticidin for 10 days to select for cells that successfully integrated the PEmax construct. These polyclonal cells were single-cell sorted into 96-well plates to grow up clonal cell lines to generate a MLH1ko-PEMax-PC9 cell line.

##### Prime editing experiment to assess editing efficiencies of 15 monoclonal lines

15 monoclonal lines were expanded, and prime editing efficiency of 12 of these lines were tested by performing an arrayed experiment in which we tested for the ability of these cells to insert a trinucleotide sequence (CTT) at the *HEK3* locus when transfected with a pegRNA that programs this insertion^22^ (**Fig. S4**). From this experiment, we chose to use Clone A6 for all further prime editing experiments. While Clones A8 and A12 exhibited higher editing efficiency (Fig. S4), these clones grew poorly and were not considered for future experiments.

#### pegRNA selection and design

All pegRNAs were designed using either the PrimeDesign^49^ web or command line interface, using default parameters. Up to four pegRNAs were designed for each individual programmed edit. Input files used for designing pegRNAs are in **Tables S3 and S7**.

#### pegRNA cloning into transient and lentiviral vectors

##### EGFR C797S T>A transiently transfected editing experiments

For the EGFR transient C797S T>A editing experiments, the following three pegRNA containing oligos (denoted pegRNA_1, pegRNA_2, and pegRNA_3) were ordered as three separate oPools from Integrated DNA Technologies (IDT):

1. pegRNA_1: CACCG**ATCACGCAGCTCATGCCCTT**GTTTTAGAGCTAGAAATAGCAAGTTAAAATAAGGCTAGTCCGTTAT CAACTTGAAAAAGTGGGACCGAGTCGGTCC**TCCAGGAGGC*T*GCCGAAGGGCATGAGCTGC**TTTT
2. pegRNA_2: CACC**GTTCCCGGACATAGTCCAGG**GTTTTAGAGCTAGAAATAGCAAGTTAAAATAAGGCTAGTCCGTTATC AACTTGAAAAAGTGGGACCGAGTCGGTCC**TTCGGC*A*GCCTCCTGGACTATGTCCGG**TTTT
3. pegRNA_3: CACCG**TGTGTTCCCGGACATAGTCC**GTTTTAGAGCTAGAAATAGCAAGTTAAAATAAGGCTAGTCCGTTAT CAACTTGAAAAAGTGGGACCGAGTCGGTCC**TTCGGC*A*GCCTCCTGGACTATGTCCGGGAA**TTTT

20 base pair spacer sequences and variable length extension sequences are in bold. Programmed single nucleotide edit are in bold and italics.

The pU6-pegRNA-GG-acceptor (Addgene plasmid #132777) was digested with BsaI-HFv2 (New England Biolabs, Cat. No. R3733S) in 10X rCutSmart Buffer at 37 degrees Celsius overnight to ensure complete digestion of the backbone. This digestion cuts out the mRFP1 cassette (821 base pairs). The linear backbone vector (2,183 base pairs in size) was gel extracted using a gel extraction kit (NEB, Cat. No. T1020S) and assembled with the pegRNA oligos (listed above) via Golden Gate assembly using the following amounts: 30 ng of linearized backbone, 1 uL of 1 uM pegRNA oligo, 0.25 uL of BsaI-HFv2 (New England Biolabs, Cat. No. R3733S), 0.5 uL of T4 DNA ligase (New England Biolabs, Cat. No. M0202S) and 1 uL of 10X T4 DNA ligase reaction buffer (New England Biolabs, Cat. No. B0202S) in a final volume of 10 uL. 1 uL of the assembly reaction was transformed into 50 uL of stable competent E. coli cells (New England Biolabs, Cat. No. C3040H) and plated on an LB agar plate containing ampicillin, and single colonies miniprepped and used for transfection(Zymo Research, Cat. No. D4208T).

##### EGFR C797S T>A arrayed and 121 epegRNA pooled lentiviral editing screens

For the EGFR lentiviral C797S T>A editing experiments, the following three pegRNA-containing oligos (denoted lenti_epegRNA_1, lenti_epegRNA_2, and lenti_epegRNA _3) were ordered as three separate oPools from Integrated DNA Technologies (IDT):

1. lenti_epegRNA_1: gctttatatatcttgtggaaaggacgaaacacc**GATCACGCAGCTCATGCCCTT**gttttagagctagaaatagcaagttaaaataaggctagt ccgttatcaacttgaaaaagtggGaccgagtcggtCc**TCCAGGAGGC*T*GCCGAAGGGCATGAGCTGC**TTGACGCGGTTCT ATCTAGTTACGCGTTAAACCAACTAGAAAtttttttNNNNNNNNggagacgaagcttggcg
2. lenti_epegRNA_2: gctttatatatcttgtggaaaggacgaaacacc**GGTTCCCGGACATAGTCCAGG**gttttagagctagaaatagcaagttaaaataaggctagt ccgttatcaacttgaaaaagtggGaccgagtcggtCc**TTCGGC*A*GCCTCCTGGACTATGTCCGG**TTGACGCGGTTCTATCT AGTTACGCGTTAAACCAACTAGAAAtttttttNNNNNNNNggagacgaagcttggcg
3. lenti_epegRNA_3: gctttatatatcttgtggaaaggacgaaacacc**GTGTGTTCCCGGACATAGTCC**gttttagagctagaaatagcaagttaaaataaggctagt ccgttatcaacttgaaaaagtggGaccgagtcggtCc**TTCGGC*A*GCCTCCTGGACTATGTCCGGGAA**TTGACGCGGTTCT ATCTAGTTACGCGTTAAACCAACTAGAAAtttttttNNNNNNNNggagacgaagcttggcg

20 base pair spacer sequences and variable length extension sequences are in bold. Programmed single nucleotide edit are in bold and italics.

The LentiGuide-Puro-P2A-EGFP (Addgene plasmid #137729) was modified to enable cloning of epegRNAs via Gibson assembly. The modified plasmid, which we term Lenti-epeg-Puro-P2A-EGFP, was used for cloning all epegRNAs. Lenti-epeg-Puro-P2A-EGFP was digested with BsmBI-v2 (New England Biolabs, Cat. No. R0739S) to create a single cut and produce a linearized backbone vector of 9007 base pairs. The three lenti_epegRNA oligos were PCR amplified with the following primers: gctttatatatcttgtggaaaggacg (forward primer) and cgccaagcttcgtctcc (reverse primer) to create double stranded DNA. The double-stranded lenti_epegRNA oligos were then assembled with the linearized Lenti-epeg-Puro-P2A-EGFP backbone using NEBuilder^®^ HiFi DNA Assembly Master Mix (New England Biolabs, Cat. No. E2621S) using the standard protocol. 1 uL of the 20 uL assembly reaction was transformed into 50 uL of stable competent E. coli cells (New England Biolabs, Cat. No. C3040H), plated on an LB agar plate containing ampicillin, and single colonies were miniprepped (Zymo Research, Cat. No. D4208T) and used used for lentivirus generation.

##### 1,220 variant lentiviral pooled editing screen

epegRNA-containing oligos were ordered as four separate sub-libraries in one single oligo pool from Twist biosciences. Sequences in this oligo pool are in **Table S2.** The four sub-libraries were cloned separately into the Lenti-epeg-Puro-P2A-EGFP vector, following the steps described above for the 121 epegRNA pool, with the only difference being that after assembly, 2 uL of each library was transformed into 50 uL of electrocompetent E. coli cells (New England Biolabs, Cat. No. C3020K) via electroporation. 990 uL of transformed cells were cultured in 50 mL of LB media + 100 ug/mL ampicillin at 31C. 10 uL (out of a total volume of 1000 uL) of the transformed E. coli cells was plated on LB agar plates containing ampicillin, and colonies were counted the next day to estimate the number of clones in each library. Transformations were performed for each library until each library had a minimum of 1000X coverage of each epegRNA. After overnight culture at 31C, the epegRNA lentiviral plasmid libraries were extracted using a Qiagen Plasmid Midi Kit (Qiagen, Cat. No. 12143). Extracted plasmid DNA from the four libraries was pooled to generate a pool with 3,825 epegRNAs and used for lentivirus generation.

#### Lentivirus generation

To generate lentivirus, HEK293T cells (ATCC, Cat. No. CRL-3216) were either plated the day before transfection at 0.7 ×10^6^ cells in a T-25 cell culture flask, or the day of transfection at 1.4 ×10^6^ cells in a T-25 cell culture flask. Virapower lentiviral packaging mix (ThermoFisher Scientific, Cat. No. K497500) was used with Lipofectamine 3000 (ThermoFisher Scientific, Cat. No. L3000001) for transfection into HEK293T cells. 1,500 uL of Opti-MEM reduced serum media (Cat. No. 31985062) was mixed with 42 uL of Lipofectamine 3000 in one tube. 13.5 ug of Virapower lentiviral packaging mix, 4.5 ug of the epegRNA lentiviral plasmid or plasmid library, and 36 uL of P3000 reagent was added to a second tube. The two tubes were combined into a single tube and incubated for 10-20 minutes at room temperature. 50% of the media was removed from the T-25 flask containing the HEK293T cells (unless the cells were plated the day of, then they were only plated in half the amount of media), and the lipid complex from the single tube was added to the cells. The cells were incubated overnight, and media harvests were taken at 24, 48, and 72 hours post-transfection. The lentivirus-containing media was mixed at a 1:4 ratio of PEG-it virus precipitation solution (System Biosciences, Cat. No. LV810A-1) to media, and refrigerated at 4C to concentrate the lentiviral particles. After 96 hours, all three harvests (24, 48, and 72 hours) were spun down at 1,500xg for 30 minutes at 4C. The lentivirus was a visible white pellet after this spin. The media above the pellet was aspirated, and the lentivirus was resuspended in 400 uL ice cold 1X PBS (Invitrogen, Cat. No. 14190-144), aliquoted into 100 uL aliquots, and frozen at -80 for later use. Each lentiviral aliquot was only thawed a single time prior to use to avoid multiple freeze-thaw cycles. For the larger scale screens, all amounts were increased in scale to make more lentivirus in larger cell culture dishes (T-75 cell culture flasks).

#### Lentivirus titration experiment in 3,825 epegRNA lentiviral pooled editing screens

A titration experiment was performed to determine lentivirus amounts to achieve the desired multiplicity of infection (MOI) for both screens. 100,000 PC-9 cells were seeded into each well of a 12-well plate in 1 mL of RPMI 1640 + L-Glutamine media (GIBCO, Cat. No. <u>11-875-093</u>). 0, 0.5, 1, 2, 4, and 8 uL of virus was added to each of 2 wells in the 12-well plate (2 replicates per lentivirus amount). 48 hours after transduction, varying amounts of GFP were observed in the different conditions (GFP is expressed off the epegRNA vector), indicating successful transduction. MOI was determined by flow cytometry (**Fig. S4**). 3 uL of virus for every 100,000 cells was the condition used for the large scale screen to achieve ∼30% GFP positivity, which represents an MOI of ∼0.35.

#### Osimertinib dose titration curves

*MLH1*ko-PEmax PC-9 cells expressing either no epegRNA (termed control cells) or an epegRNA coding (lenti_epegRNA_2) for an EGFR C797S mutation (termed EGFR^C797S^ cells) were seeded in duplicate in 12-well dishes at a density of 50,000 cells per well (approximately 14,300 cells/cm^2^) in a total volume of 2 mL. Cells were treated with either vehicle (DMSO, Sigma-Aldrich Cat. No. D2650) at a concentration of 500 nM or osimertinib (AZD9291, SelleckChem Cat. No. S7297) at a concentration of 100, 300, or 500 nM. From this point, cells were passaged every 3 days and replated at a density of 50,000 cells per well in the appropriate concentration of either vehicle or osimertinib. To measure growth rate and cumulative population doublings of control versus EGFR^C797S^ cells in vehicle and osimertinib, cells were counted at each passage using a Vi-CELL XR analyzer (Beckman Coulter). The 300 nM osimertinib dose, which led to an appreciable growth rate difference between control and EGFR^C797S^ cells was used for all further experiments (**Fig. S2**).

#### pegRNA transient transfection or lentiviral transduction into PC-9 cells

##### Transient transfections

For the EGFR C797S T>A proof of concept experiment, the SF Cell Line 4D-Nucleofector X Kit S (Lonza, Cat. No. V4XC-2032) was used to transfect the three pegRNAs and one no DNA control. 2.5 ug of DNA (the pegRNA plasmid and the pCMV-PE2 plasmid (Addgene #132775) at a 1:2 ratio by mass of pegRNA plasmid:PE2), was transfected into 100,000 PC-9 cells that went into four wells of a 12-well plate. 2 days later, 400 nM osimertinib was added to the cells for all conditions (no drug, and the three separate pegRNAs). 24 days later, cells were harvested and the EGFR C797 locus was amplified and sequenced.

##### Lentiviral transductions

A ratio of 3 uL of lentivirus was added for every 100,000 cells transduced. Cell culture amounts were scaled as necessary to transduce cells at 2,000X coverage of epegRNAs. One day after transduction, media was aspirated and replenished with fresh media. Two days after lentiviral transduction, 2 ug/mL puromycin (GIBCO/Thermo Fisher Scientific, Cat. No. <u>A11138</u>03) was added to the cells for 4 days to select for successfully transduced cells. The cells were replenished with fresh media with 2 ug/mL puromycin each day during these 4 days. For the arrayed *EGFR* C797S T>A experiment, after puromycin selection, we transiently transfected cells with PB-EFS-PEmax twice on two consecutive days. On the second transfection day, cells were treated with 200 nM osimertinib. Eight days later, cells were harvested and the *EGFR* locus was amplified and sequenced. For the 3,825 epegRNA lentiviral pooled editing screens, two parallel plates were cultured without puromycin selection to enable a FACS analysis one week later to determine the MOI of this screen (**Fig. S5**). After 4 days of puromycin selection, 300 nM of drug (for all other lentiviral editing experiments) was added to each treatment condition (CH7233163, osimertinib, or sunvozertinib), and the same volume of DMSO was added to the no treatment control condition. The day that the drug treatments were added marks Day 0 of the screens.

#### Drug treatments used in screens

Cells were treated with DMSO (SigmaAldrich, Cat. No. D2438) or 100, 300, or 500 nM of CH7233163 (Selleckchem, Cat. No. S9711), osimertinib (Selleckchem, Cat. No. S7297), or sunvozertinib (Selleckchem, Cat. No. E0368), depending on the screen. Drugs were initially diluted to 1 uM in DMSO, and all further dilutions were done in PBS (Invitrogen, Cat. No. 14190-144) to reach the desired concentrations.

#### Cell harvests and genomic DNA extraction for the 3,825 epegRNA lentiviral pooled editing screens

Cells were harvested on day 8 for the lentiviral EGFR C797S T>A proof of concept experiment. Cells were harvested on days 3, 7, 11, 15, and 19 for both of the large scale screens. Cells were harvested such that a minimum of 500X coverage of epegRNAs was retained in the remaining culture. At each harvest, fresh media and drug (DMSO, CH7233163, osimertinib, or sunvozertinib) was added to the passaged cells. The harvested cells were spun down at 400 x g for 5 minutes, aspirated, and cell pellets were stored at -20C. The Puregene Cell Kit (8×10^8^) (Qiagen, Cat. No. 158767) was used for all genomic DNA (gDNA) extractions for the large scale screens using the standard protocol, with a single modification to add 2 uL of GlycoBlue coprecipitant (ThermoFisher Scientific, Cat. No. AM9515) to the DNA pellet.

#### pegRNA amplicon amplification and sequencing

##### Amplification of the EGFR C797 locus

For the EGFR C797S T>A proof of concept transient and lentiviral editing experiments, the EGFR locus was PCR amplified with the following primers: GCGTCAGATGTGTATAAGAGACAG**CATCTGCCTCACCTCCACCGTG** (forward primer) and GTGACTGGAGTTCAGACGTGTGCTCTTCCGATCT**ACCAGTTGAGCAGGTACTGGGAGC** (reverse primer). The sequences in bold are the locus-binding part of the primer, and the sequences not in bold contain Nextera and Truseq adapter sequences. KAPA2G Robust HotStart ReadyMix (Roche Diagnostics, Cat. No. KK5702) was used for amplification using the recommended protocol with a 60C annealing temperature and a 30 second extension time. The product of this PCR was cleaned with a 1X AMpure XP bead cleanup (Beckman Coulter, Cat. No. A63880), and eluted in 20 uL of water. 1 uL of cleaned PCR product was used for a second PCR which adds the Illumina P5 and P7 adapter sequences and sample-specific indices. This second PCR was done using the following primers: AATGATACGGCGACCACCGAGATCTACACNNNNNNNNNNTCGTCGGCAGCGTCAGATGTGTATAAGAGACAG (forward primer) and CAAGCAGAAGACGGCATACGAGATNNNNNNNNNNGTGACTGGAGTTCAGACGTGTGCTCTTCCGATCT (reverse primer). 10N sequences denote unique indices for each sample. KAPA2G Robust HotStart ReadyMix (Roche Diagnostics, Cat. No. KK5702) was used for amplification using the recommended protocol with a 60C annealing temperature and a 30 second extension time for this second PCR. The product of this PCR was cleaned with a 1X AMpure XP bead cleanup (Beckman Coulter, Cat. No. A63880), and eluted in 20 uL of water. Amplicon concentration and size was determined using the Qubit (ThermoFisher Scientific) and a Tapestation (Agilent) instrument.

##### Amplification of the integrated genomic DNA pegRNA construct

For the small (121 epegRNAs) and large (3,825 epegRNAs) scale screens, the genomically integrated epegRNAs were amplified via PCR. This is a two-step PCR, with the first PCR amplifying the epegRNA construct and adding partial Illumina read adapter sequences, and with the second PCR adding the Illumina P5 and P7 adapter sequences and sample-specific indices. The first PCR uses the following primers: GCGTCAGATGTGTATAAGAGACAG**cttgtggaaaGGACGAAACACC** (forward primer) and ACGTGTGCTCTTCCGATCT**tctcaagatctagttacgccaagc** (reverse primer). The sequences in bold are the locus-binding part of the primer, and the sequences not in bold contain Nextera and Truseq adapter sequences. KAPA2G Robust HotStart ReadyMix (Roche Diagnostics, Cat. No. KK5702) was used for the 121 epegRNA small scale screen and KAPA HiFi HotStart ReadyMix (Roche Diagnostics, Cat. No. KK2602) was used for amplification in the large scale screens using the recommended protocol with a 65C annealing temperature for the KAPA2G Robust PCR and the KAPA HiFi PCR, and a 30 second extension time for both protocols. The product of this PCR was cleaned with a 1X AMpure XP bead cleanup (Beckman Coulter, Cat. No. A63880), and eluted in 20 uL of water. 5 uL of cleaned PCR product was used for a second PCR which adds the Illumina P5 and P7 adapter sequences and sample-specific indices. This second PCR was done using the following primers: AATGATACGGCGACCACCGAGATCTACACNNNNNNNNNNTCGTCGGCAGCGTCAGATGTGTATAAGAGACAG (forward primer) and CAAGCAGAAGACGGCATACGAGATNNNNNNNNNNGTGACTGGAGTTCAGACGTGTGCTCTTCCGATCT (reverse primer). 10N sequences denote unique indices for each sample. KAPA2G Robust HotStart ReadyMix (Roche Diagnostics, Cat. No. KK5702) was used for the 121 epegRNA small scale screen and KAPA HiFi HotStart ReadyMix (Roche Diagnostics, Cat. No. KK2602) was used for amplification in the large scale screen using the recommended protocol with a 65C annealing temperature for the KAPA2G Robust PCR and the KAPA HiFi PCR, and a 30 second extension time was used for this second PCR for both protocols. The product of this PCR was cleaned with a 1X AMpure XP bead cleanup (Beckman Coulter, Cat. No. A63880), and eluted in 20 uL of water. Amplicon concentration and size was determined using the Qubit (ThermoFisher Scientific) and a Tapestation (Agilent) instrument.

##### Amplification of endogenous loci targeted with prime editing gRNAs

For the 121 epegRNA lentiviral pooled experiment, the endogenous locus of each target was PCR amplified and sequenced. Each locus was amplified with locus-specific primers which are listed in **Table S5**. KAPA2G Robust HotStart ReadyMix (Roche Diagnostics, Cat. No. KK5702) was used for amplification using the recommended protocol with a 60C annealing temperature and a 30 second extension time. The product of this PCR was cleaned with a 1X AMpure XP bead cleanup (Beckman Coulter, Cat. No. A63880), and eluted in 20 uL of water. 1 uL of cleaned PCR product was used for a second PCR which adds the Illumina P5 and P7 adapter sequences and sample-specific indices. This second PCR was done using the following primers: AATGATACGGCGACCACCGAGATCTACACNNNNNNNNNNTCGTCGGCAGCGTCAGATGTGTATAAGAGACAG (forward primer) and CAAGCAGAAGACGGCATACGAGATNNNNNNNNNNGTGACTGGAGTTCAGACGTGTGCTCTTCCGATCT (reverse primer). 10N sequences denote unique indices for each sample. KAPA2G Robust HotStart ReadyMix (Roche Diagnostics, Cat. No. KK5702) was used for amplification using the recommended protocol with a 60C annealing temperature and a 30 second extension time for this second PCR. The product of this PCR was cleaned with a 1X AMpure XP bead cleanup (Beckman Coulter, Cat. No. A63880), and eluted in 20 uL of water. Amplicon concentration and size was determined using the Qubit (ThermoFisher Scientific) and a Tapestation (Agilent) instrument.

#### Sequencing of prime editing screening libraries

Final libraries were sequenced on either a Miseq 300 cycle kit or a NextSeq 2000 P3 200 cycle kit with standard Illumina Nextera and Truseq adapter sequences. The pegRNA spacer was sequenced on Read 1, and the prime editing gRNA extension sequence was sequenced on Read 2. 10 bp index sequences were sequenced with 10 cycle index 1 and index 2 reads.

#### Raw data processing and quality control filtering of sequencing data

##### Fastq file generation, pegRNA read counting, and z-score calculation

Bcl2fastq version 2.20 was used to generate fastq files using default parameters. Fastq files were then input into count_reads.py via a Snakemake pipeline to count the occurrence of each pegRNA in each day, drug treatment, and replicate condition. A pseudocount of 1 was added to all pegRNA read counts, and then read counts were log2 normalized by condition to normalize for variable sequencing depths across samples. This log2 normalized count matrix was used to calculate the log2 fold change of each pegRNA between each treatment condition and the DMSO control treatment condition. Finally, a z-score was calculated for each pegRNA, which is equal to:

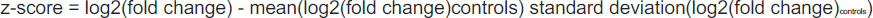

Controls are pegRNAs that encode for synonymous edits.

##### DESeq2 for differential pegRNA abundance analysis

DESeq2 was used to identify differential pegRNA abundances in the large screen that programmed 1,220 variants with 3,825 epegRNAs in ten oncogenes. The count matrix that was used as an input to DESeq2 was generated from the raw read counts (i.e. not log2 normalized) from the sequencing data. DESeq2 was run for each of the three drug screens separately (osimertinib, sunvozertinib, and CH7233163), using the likelihood ratio test (LRT) for statistical testing. The full model includes the variables time, drug treatment, and drug:time (the latter interaction term tests for the effect of drug as a result of time), and the reduced model contains only the variable time. The likelihood ratio between the full and reduced model tests if the increased likelihood of the data using the full model is more than expected (i.e. more than 1) and if the parameters in the full model fit the data better than the parameters in the reduced model. DESeq2 was run using default parameters, except for when running the DESeq() function, the parameter minReplicatesForReplace=Inf was used, and in the results() function, the additional parameters cooksCutoff=FALSE and independentFiltering=FALSE were used. These additional parameters reduce the filtering of the data to include outlier data (i.e. data with higher read counts). This data was not filtered out because the high read counts are coming from variants with high resistance phenotypes (i.e. EGFR C797S and KRAS G12 variants). Because DESeq2 is typically used for gene expression analyses, these parameters were appropriately changed for this drug resistance screening data type.

All further downstream analyses were done using custom Python and R scripts, which will soon be accessible on the following website: https://krishna.gs.washington.edu/content/members/multiplex_PE_screening/public/.

## Notes

### Competing Interest Statement

Author Shendure is a scientific advisory board member, consultant and/or co-founder of Cajal Neuroscience, Guardant Health, Maze Therapeutics, Camp4 Therapeutics, Phase Genomics, Adaptive Biotechnologies, Scale Biosciences, Sixth Street Capital, Pacific Biosciences, and Prime Medicine. All other authors declare no competing interests.

